# Association of Peripheral Blood Pressure with Grey Matter Volume in 19- to 40-Year-Old Adults

**DOI:** 10.1101/239160

**Authors:** H. Lina Schaare, Shahrzad Kharabian Masouleh, Frauke Beyer, Deniz Kumral, Marie Uhlig, Janis D. Reinelt, Andrea M.F. Reiter, Leonie Lampe, Anahit Babayan, Miray Erbey, Josefin Roebbig, Matthias L. Schroeter, Hadas Okon-Singer, Karsten Müller, Natacha Mendes, Daniel S. Margulies, A. Veronica Witte, Michael Gaebler, Arno Villringer

**Affiliations:** Department of Neurology, Max Planck Institute for Human Cognitive and Brain Sciences, Leipzig, Germany; International Max Planck Research School NeuroCom, Leipzig, Germany; MindBrainBody Institute at Berlin School of Mind and Brain, Charité & Humboldt Universität zu Berlin, Germany; Lifespan Developmental Neuroscience, Technische Universität Dresden, Germany; Leipzig Research Centre for Civilization Diseases (LIFE), University of Leipzig, Germany; Max Planck Research Group for Neuroanatomy & Connectivity, Max Planck Institute for Human Cognitive and Brain Sciences, Leipzig, Germany; Nuclear Magnetic Resonance Group, Max Planck Institute for Human Cognitive and Brain Sciences, Leipzig, Germany; Department of Psychology, University of Haifa, Israel; Clinic for Cognitive Neurology, University of Leipzig, Germany; Collaborative Research Centre 1052 ‘Obesity Mechanisms’, Subproject A1, Faculty of Medicine, University of Leipzig, Germany; Center for Stroke Research Berlin, Charité – Universitätsmedizin Berlin, Germany

**Keywords:** Stroke prevention [12], Vascular dementia [32], All Cerebrovascular disease/Stroke [2], Risk factors in epidemiology [59], MRI [120]

## Abstract

**Objective:** To test whether elevated blood pressure (BP) relates to grey matter volume (GMV) changes in young adults who had not previously been diagnosed as hypertensive (systolic BP (SBP)/diastolic BP (DBP)≥140/90 mmHg).

**Methods:** We associated BP with GMV from structural 3 Tesla T1-weighted MRI of 423 healthy adults between 19-40 years (mean age=27.7±5.3 years, 177 women, SBP/DBP=123.2/73.4±12.2/8.5 mmHg). Data originated from four previously unpublished cross-sectional studies conducted in Leipzig, Germany. We performed voxel-based morphometry on each study separately and combined results in image-based meta-analyses (IBMA) to assess cumulative effects across studies. Resting BP was assigned to one of four categories: (1) SBP<120 and DBP<80 mmHg, (2) SBP 120-129 or DBP 80-84 mmHg, (3) SBP 130-139 or DBP 85-89 mmHg, (4) SBP≥140 or DBP≥90 mmHg.

**Results:** IBMA yielded: (a) lower regional GMV was correlated with higher peripheral BP; (b) lower GMV with higher BP when comparing individuals in sub-hypertensive categories 3 and 2, respectively, to those in category 1; (c) lower BP-related GMV was found in regions including hippocampus, amygdala, thalamus, frontal and parietal structures (e.g. precuneus).

**Conclusions:** BP≥120/80 mmHg was associated with lower GMV in regions that have previously been related to GM decline in older individuals with manifest hypertension. Our study shows that BP-associated GM alterations emerge continuously across the range of BP and earlier in adulthood than previously assumed. This suggests that treating hypertension or maintaining lower BP in early adulthood might be essential for preventing the pathophysiological cascade of asymptomatic cerebrovascular disease to symptomatic end-organ damage, such as stroke or dementia.

## Introduction

Hypertension (HTN) is highly prevalent and the leading single risk factor for global disease burden and overall health loss^1,2^. The risk for insidious brain damage and symptomatic cerebrovascular disease (CVD, e.g. stroke and vascular dementia) multiplies with manifestation of HTN^3^. Midlife HTN is a major risk factor for late-life cognitive decline and has been associated with risk for dementia, including late-onset Alzheimer’s disease (AD)^3–5^.

Importantly, HTN is also related to sub-clinical functional^6,7^ and structural^5–14^ brain changes, or *asymptomatic* CVD, including brain volume reductions in the medial temporal and frontal lobes^5,6,9–11,15^. Hippocampal volumes, in particular, have been consistently associated with HTN-related reductions^5,9,10,15^. Furthermore, computational anatomy has been employed to detect subtle cerebral changes, such as microstructural white matter (WM) alterations ^13^ or reductions in regional grey matter^5,11^, in middle-aged and older adults with elevated blood pressure (BP).

Recent statements suggest that symptomatic clinical disease, resulting from elevated BP, could be prevented by avoiding primary BP elevations and sub-clinical target organ damage (including brain damage) in early adulthood and middle-age^3,16,17^. However, effects of elevated BP on adult brains before the age of 40 are unclear. Preliminary evidence from 32 young, normotensive adults, showed that BP-reactivity correlated with lower amygdala volume^18^.

This study aimed to investigate if subtle structural brain changes occur in early adulthood (<40 years) at sub-hypertensive BP levels. We hypothesized that higher BP would relate to lower regional grey matter volume (GMV) and that this would predominantly affect frontal and medial temporal lobes, including amygdala and hippocampus.

## Methods

We applied voxel-based morphometry^19,20^ (VBM) to four previously unpublished independent datasets including young adults aged between 19-40 years without previous diagnosis of HTN or any other severe, chronic or acute disease. Results from each dataset were combined in image-based meta-analyses (IBMA) for well-powered, cumulative evaluation of findings across study differences (i.e. recruitment procedure, inclusion criteria and data acquisition, Figure 1, data available from bioRxiv (Supplementary Table 1) https://doi.org/10.1101/239160).

**Figure 1.**
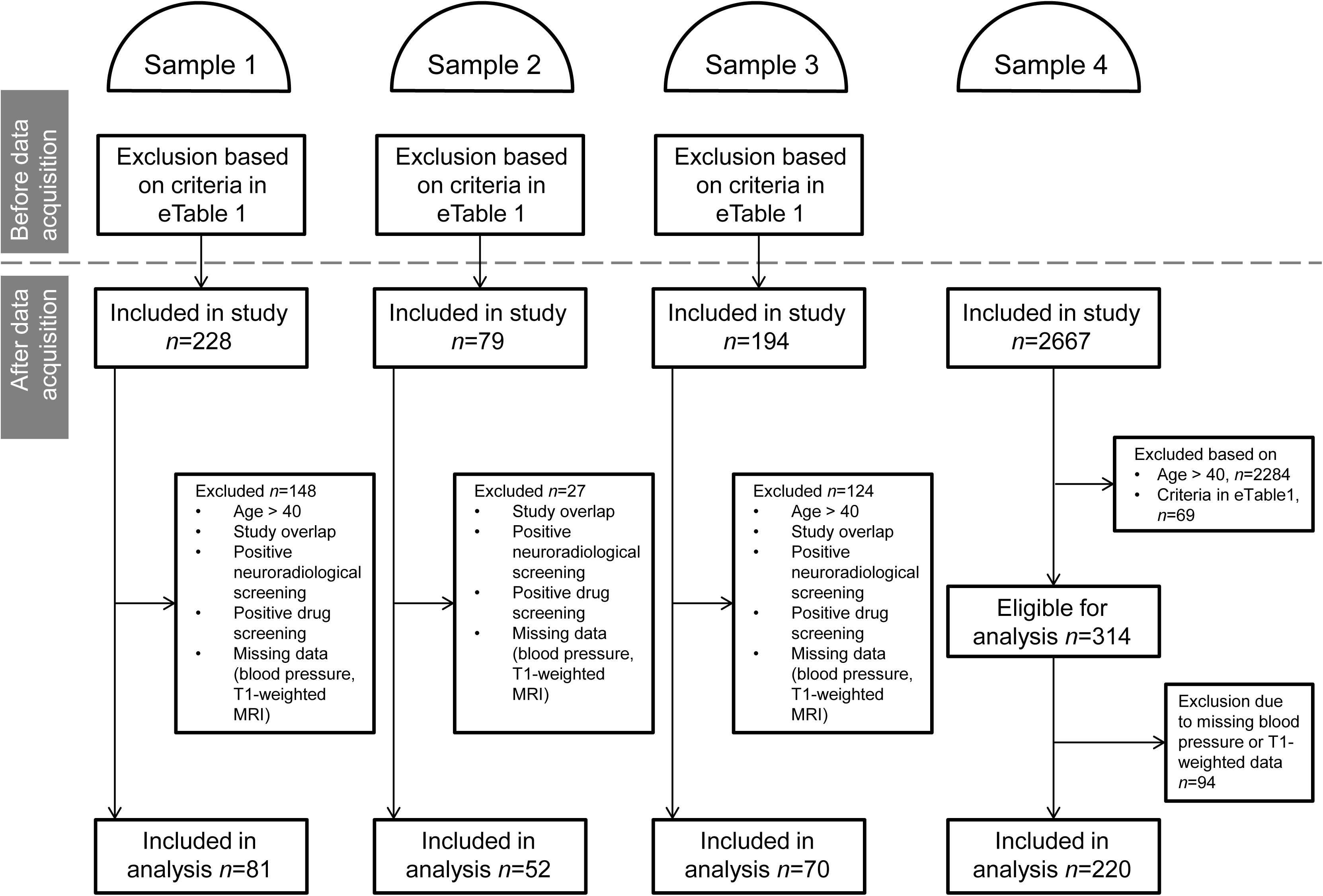
Flow chart with inclusion procedure for the four samples. Sample 1: Leipzig Study for Mind-Body-Emotion Interactions. Sample 2: Neural Consequences of Stress Study. 3: Neuroanatomy and Connectivity Protocol. 4: Leipzig Research Centre for Civilization Diseases (MRI-subcohort).

### Participants

We included cross-sectional data of 423 young participants from four samples. The samples were drawn from larger studies that were conducted in Leipzig, Germany, between 2010-2015: 1. Leipzig Study for Mind-Body-Emotion Interactions (Babayan et al., under review), 2. Neural Consequences of Stress Study (Reinelt et al., in preparation), 3. Neuroanatomy and Connectivity Protocol^21^, 4. Leipzig Research Centre for Civilization Diseases (LIFE)^22^.

The objective of study 1 was to cross-sectionally investigate mind-brain-body-emotion interactions in a younger (20-35 years) and an older (59-77 years) group of 228 healthy volunteers. Study 2 aimed to investigate neural correlates of acute psychosocial stress in 79 young (18-35 years), healthy, non-smoking men. The study protocol for the baseline assessment of participants in study 2 was adapted from the protocol in study 1. In study 3, 194 healthy volunteers between 20-75 years of age participated in one session of MRI and completed an extensive assessment of cognitive and personality measures. This dataset aimed to relate intrinsic functional brain connectivity with cognitive faculties, self-generated mental experience, and personality features. Together, studies 1-3 constitute the MPI-Leipzig Mind-Brain-Body database. Study 4 (LIFE-Study) is a population-based dataset in the city of Leipzig, Germany, with the objective to investigate the development of major modern diseases. Overall, 10000 participants were randomly drawn from the local population of whom 2667 underwent MRI and detailed screening. With dementia being one of the key scientific topics in this study, most participants in the MRI-subcohort were adults above the age of 60 years. The exact inclusion procedure and numbers for the current investigation is depicted in Figure 1. Inclusion criteria for our study were age between 19-40 years, availability of high-resolution structural T1-weighted MRI and ≥1 BP measurements. Participants were excluded in case of previously diagnosed HTN, intake of antihypertensive drugs or severe diseases (data available from bioRxiv (Supplementary Table 1) https://doi.org/10.1101/239160).

#### Standard Protocol Approvals, Registrations, and Patient Consents

The studies were in agreement with the Declaration of Helsinki and approved by the ethics committee of the medical faculty at the University of Leipzig, Germany (ethics reference numbers study 1: 154/13-ff, study 2: 385/14-ff, study 3: 097/15-ff, study 4: 263-2009-14122009). Before entering the studies, participants gave written informed consent.

### Blood pressure measurements

Systolic (SBP) and diastolic blood pressure (DBP) were measured at varying times of day using an automatic oscillometric blood pressure monitor (OMRON M500 (samples 1-3), 705IT (sample 4), OMRON Medizintechnik, Mannheim, Germany) after a seated resting period of 5 min. In sample 1, three measurements were taken from participants’ left arms on three separate occasions within two weeks. In sample 2, two measurements were taken from participants’ left arms on two separate occasions on the same day. In sample 3, blood pressure was measured once before participants underwent MRI. In sample 4, the procedure consisted of three consecutive blood pressure measurements, taken from the right arm in intervals of 3 minutes. In each sample, all available measurements per participant were averaged to one systolic and one diastolic blood pressure value. These averages were used for classification of BP.

### Neuroimaging

MRI was performed at the same 3 Tesla MAGNETOM Verio Scanner (Siemens, Erlangen, Germany) for all studies with a 32-channel head coil. Whole-brain 3-dimensional T1-weighted volumes with a resolution of 1 mm isotropic were acquired for the assessment of brain structure. T1-weighted images in sample 4 were acquired with a standard MPRAGE protocol (inversion time TI=900 ms, repetition time TR=2300 ms, echo time TE=2.98 ms, flip angle FA=9°, field of view FOV=256×240×176 mm^3^, voxel size=1×1×1 mm^3^), while T1-weighted images in samples 1-3 resulted from an MP2RAGE protocol (TI1=700 ms, TI2=2500 ms, TR=5000 ms, TE=2.92 ms, FA1=4°, FA2=5°, FOV=256×240×176 mm ^3^, voxel size=1×1×1 mm^3^). Grey and white matter contrast are comparable for the two sequence protocols^1,2^, but additional preprocessing steps were performed for MP2RAGE T1-weighted images (data available from bioRxiv (Additional Methods) https://doi.org/10.1101/239160). Fluid-attenuated inversion recovery (FLAIR) images were acquired in all samples for radiological examination for incidental findings and for Fazekas scale ratings for white matter hyperintensities (WMH, Tables 1 and 2).

**Table 1.** Characteristics by blood pressure category. Characteristics of the total sample by blood pressure categories. Column *p* specifies significant results of comparisons between blood pressure categories: empty cells = *p*>0.05. Column *Pairwise comparisons* specifies significant post-hoc comparisons for: 2=category 1 vs. 2, 3=category 1 vs. 3, 4=category 1 vs. 4. ***= *p*<0.001, **=*p*<0.01, *=*p*<0.05. Definition of blood pressure categories: *category 1* (SBP<120 mmHg and DBP<80 mmHg), *category 2* (SBP 120-129 mmHg or DBP 80-84 mmHg), *category 3* (SBP 130-139 mmHg or DBP 85-89 mmHg) and *category 4* (SBP≥140 mmHg or DBP≥90 mmHg).

|  | Total |  | Category 1 |  | Category 2 |  | Category 3 |  | Category 4 |  | *p* | Pairwise comparisons |
| --- | --- | --- | --- | --- | --- | --- | --- | --- | --- | --- | --- | --- |
| **n (%)** | 423 (100%) |  | 175 (41%) |  | 121 (29%) |  | 80 (19%) |  | 47 (11%) |  |  |  |
| Women [n (%)] | 177 (42%) |  | 117 (67%) |  | 40 (33%) |  | 11 (14%) |  | 9 (19%) |  | *** | 2***, 3***, 4*** |
| **Age (mean years, SD)** | 27.66 | 5.27 | 27.61 | 5.53 | 27.30 | 4.95 | 28.01 | 5.24 | 28.21 | 5.23 |  |  |
| Range (min-max years) | 19-40 |  | 19-40 |  | 20-40 |  | 20-40 |  | 20-39 |  |  |  |
| **Systolic Blood Pressure (mean mmHg, SD)** | 123.2 | 12.19 | 111.91 | 5.44 | 123.99 | 3.62 | 134.57 | 3.48 | 143.56 | 7.76 | *** | 2***, 3***, 4*** |
| **Diastolic Blood Pressure (mean mmHg, SD)** | 73.38 | 8.49 | 67.67 | 5.81 | 73.64 | 5.77 | 78.79 | 6.46 | 84.75 | 8.26 | *** | 2***, 3***, 4*** |
| **Body Mass Index (mean kg/m<sup>2</sup>, SD)** | 23.48 | 3.25 | 22.60 | 2.74 | 23.45 | 3.23 | 24.22 | 3.42 | 25.59 | 3.62 | *** | 2*, 3**, 4*** |
| Missing values [n (%)] | 13 (3%) |  | 5 (3%) |  | 4 (3%) |  | 1 (1%) |  | 3 (6%) |  |  |  |
| Range (min-max kg/m<sup>2</sup>) | 17.96-36.93 |  |  |  |  |  |  |  |  |  |  |  |
| **Smoking status [n (%)]** |  |  |  |  |  |  |  |  |  |  |  |  |
| non-smoker | 273 (65%) |  | 113 (65%) |  | 78 (64%) |  | 53 (66%) |  | 29 (62%) |  |  |  |
| occasional smoker | 57 (13%) |  | 23 (13%) |  | 17 (14%) |  | 13 (16%) |  | 4 (9%) |  |  |  |
| smoker | 73 (17%) |  | 29 (17%) |  | 21 (17%) |  | 13 (16%) |  | 10 (21%) |  |  |  |
| missing values | 20 (5%) |  | 10 (6%) |  | 5 (4%) |  | 1 (1%) |  | 4 (9%) |  |  |  |
| **Fazekas score [n (%)]** |  |  |  |  |  |  |  |  |  |  |  |  |
| 0 | 303 (72%) |  | 123 (70%) |  | 85 (70%) |  | 59 (74%) |  | 36 (77%) |  |  |  |
| 1 | 72 (17%) |  | 39 (22%) |  | 16 (13%) |  | 10 (13%) |  | 7 (15%) |  |  |  |
| 2 | 0 (0%) |  | 0 (0%) |  | 0 (0%) |  | 0 (0%) |  | 0 (0%) |  |  |  |
| 3 | 0 (0%) |  | 0 (0%) |  | 0 (0%) |  | 0 (0%) |  | 0 (0%) |  |  |  |
| Missing values | 48 (11%) |  | 13 (7%) |  | 20 (17%) |  | 11 (14%) |  | 4 (9%) |  |  |  |
| **Total intracranial volume (mean ml, SD)** | 1450.05 | 137.39 | 1400.49 | 127.47 | 1457.43 | 142.07 | 1508.05 | 116.23 | 1516.90 | 126.29 |  |  |
| **Grey matter volume (mean ml, SD)** | 777.41 | 88.69 | 748.55 | 79.50 | 784.29 | 90.41 | 809.72 | 87.36 | 812.16 | 86.47 |  |  |
| **White matter volume (mean ml, SD)** | 449.79 | 55.74 | 435.84 | 53.56 | 452.75 | 55.93 | 464.19 | 53.34 | 469.58 | 55.52 |  |  |
| **Cerebrospinal fluid volume (mean ml, SD)** | 222.86 | 56.60 | 216.09 | 54.57 | 220.40 | 59.53 | 234.14 | 54.14 | 235.16 | 57.34 |  |  |
| **Hippocampal volume, left (mean ml, SD)** | 3.90 | 0.45 | 3.77 | 0.41 | 3.93 | 0.46 | 4.06 | 0.43 | 4.09 | 0.43 |  |  |
| **Hippocampal volume, right (mean ml, SD)** | 3.97 | 0.43 | 3.83 | 0.40 | 4.00 | 0.44 | 4.12 | 0.40 | 4.14 | 0.39 |  |  |
| **Amygdalar volume, left (ml)** | 1.68 | 0.19 | 1.62 | 0.18 | 1.68 | 0.19 | 1.75 | 0.18 | 1.75 | 0.18 |  |  |
| **Amygdalar volume, right (ml)** | 1.50 | 0.16 | 1.45 | 0.15 | 1.51 | 0.17 | 1.56 | 0.16 | 1.57 | 0.15 |  |  |

**Table 2.** Characteristics by sample. Characteristics of the study samples. Column *p* specifies significant results of comparisons between samples: empty cells = *p*>0.05. Column *Pairwise comparisons* specifies significant post-hoc comparisons between samples: ***= *p*<0.001, **=*p*<0.01, *=*p*<0.05.

|  | Total |  | Sample 1 |  | Sample 2 |  | Sample 3 |  | Sample 4 |  | *p* | Pairwise comparisons |
| --- | --- | --- | --- | --- | --- | --- | --- | --- | --- | --- | --- | --- |
| **n** | 423 |  | 81 |  | 52 |  | 70 |  | 220 |  |  |  |
| Women [n (%)] | 177 (42%) |  | 37 (46%) |  | 0 (0%) |  | 43 (61%) |  | 97 (44%) |  | *** | 1 vs. 2***, 2 vs. 3***, 2 vs. 4***, 3 vs. 4* |
| **Age (mean years, SD)** | 27.66 | 5.27 | 24.36 | 3.07 | 25.77 | 2.44 | 26.54 | 4.82 | 29.69 | 5.65 | *** | 1 vs. 3*, 1 vs. 4***, 2 vs. 4***, 3 vs. 4*** |
| Range (min-max years) | 19-40 |  | 20-35 |  | 21-31 |  | 20-40 |  | 19-40 |  |  |  |
| **Systolic Blood Pressure (mean mmHg, SD)** | 123.2 | 12.19 | 121.8 | 12.02 | 128.4 | 9.80 | 128.4 | 12.16 | 120.8 | 11.95 | *** | 1 vs. 2**, 1 vs. 3***, 2 vs. 4***, 3 vs. 4*** |
| **Diastolic Blood Pressure (mean mmHg, SD)** | 73.38 | 8.49 | 73.27 | 6.77 | 75.68 | 8.03 | 79.91 | 7.65 | 70.80 | 8.20 | *** | 1 vs. 3***, 1 vs. 4*, 2 vs. 3**, 2 vs. 4***, 3 vs. 4*** |
| **Blood Pressure Category [n (%)]** |  |  |  |  |  |  |  |  |  |  | *** | 1 vs. 2*, 1 vs. 3**, 2 vs. 4***, 3 vs. 4*** |
| Category 1 (SBP < 120 mmHg and DBP < 80 mmHg) | 175 (41%) |  | 38 (47%) |  | 10 (19%) |  | 15 (21%) |  | 112 (51%) |  |  |  |
| Category 2 (SBP 120-129 mmHg or DBP 80-84 mmHg) | 121 (29%) |  | 23 (28%) |  | 21 (40%) |  | 19 (27%) |  | 58 (26%) |  |  |  |
| Category 3 (SBP 130-139 mmHg or DBP 85-89 mmHg) | 80 (19%) |  | 13 (16%) |  | 12 (23%) |  | 21 (30%) |  | 34 (15%) |  |  |  |
| Category 4 (SBP ≥ 140 mmHg or DBP ≥ 90 mmHg) | 47 (11%) |  | 7 (9%) |  | 9 (17%) |  | 15 (21%) |  | 16 (7%) |  |  |  |
| **Body Mass Index (mean kg/m², SD)** | 23.48 | 3.25 | 23.14 | 3.06 | 23.02 | 2.47 | 23.17 | 3.58 | 23.79 | 3.38 |  |  |
| Missing values [n (%)] | 13 (3%) |  | 0 (0%) |  | 0 (0%) |  | 12 (17%) |  | 1 (0%) |  |  |  |
| Range (min-max kg/m²) | 17.96-36.93 |  | 18.0-34.5 |  | 17.96-28.85 |  | 18.1-36.88 |  | 18.55-36.93 |  |  |  |
| **Smoking status [n (%)]** |  |  |  |  |  |  |  |  |  |  | *** | 1 vs. 2***, 1 vs. 4***, 2 vs. 3***, 2 vs. 4***, 3 vs. 4* |
| non-smoker | 273 (65%) |  | 57 (70%) |  | 52 (100%) |  | 40 (57%) |  | 124 (56%) |  |  |  |
| occasional smoker | 57 (13%) |  | 16 (20%) |  | 0 (0%) |  | 11 (16%) |  | 30 (14%) |  |  |  |
| smoker | 73 (17%) |  | 5 (6%) |  | 0 (0%) |  | 6 (9%) |  | 62 (28%) |  |  |  |
| missing values | 20 (5%) |  | 3 (4%) |  | 0 (0%) |  | 13 (19%) |  | 4 (2%) |  |  |  |
| **Fazekas score [n (%)]** |  |  |  |  |  |  |  |  |  |  |  |  |
| 0 | 303 (72%) |  | 60 (74%) |  | 15 (29%) |  | 49 (70%) |  | 179 (81%) |  |  |  |
| 1 | 72 (17%) |  | 14 (17%) |  | 4 (8%) |  | 16 (23%) |  | 38 (17%) |  |  |  |
| 2 | 0 (0%) |  | 0 (0%) |  | 0 (0%) |  | 0 (0%) |  | 0 (0%) |  |  |  |
| 3 | 0 (0%) |  | 0 (0%) |  | 0 (0%) |  | 0 (0%) |  | 0 (0%) |  |  |  |
| Missing values | 48 (11%) |  | 7 (9%) |  | 33 (63%) |  | 5 (7%) |  | 3 (1%) |  |  |  |
| **Total intracranial volume (mean ml, SD)** | 1450.05 | 137.39 | 1448.41 | 138.07 | 1553.45 | 100.05 | 1424.99 | 127.14 | 1434.19 | 137.82 | *** | 1 vs. 2***, 2 vs. 3***, 2 vs. 4*** |
| **Grey matter volume (mean ml, SD)** | 777.41 | 88.69 | 829.38 | 74.94 | 880.24 | 59.62 | 805.25 | 64.19 | 725.11 | 66.89 | *** | 1 vs. 2***, 1 vs. 3*, 1 vs. 4***, 2 vs. 3***, 2 vs. 4***, 3 vs. 4*** |
| **White matter volume (mean ml, SD)** | 449.79 | 55.74 | 430.85 | 49.28 | 467.25 | 40.38 | 420.51 | 46.23 | 461.95 | 58.48 | *** | 1 vs. 2***, 1 vs. 4***, 2 vs. 3***, 3 vs. 4*** |
| **Cerebrospinal fluid volume (mean ml, SD)** | 222.86 | 56.60 | 188.19 | 42.88 | 205.97 | 36.83 | 199.22 | 48.70 | 247.13 | 56.18 | *** | 1 vs. 4***, 2 vs. 4***, 3 vs. 4*** |
| Hippocampal volume, left (mean ml, SD) | 3.90 | 0.45 | 4.17 | 0.39 | 4.42 | 0.33 | 4.03 | 0.35 | 3.65 | 0.32 | *** | 1 vs. 2***, 1 vs. 3*, 1 vs. 4***, 2 vs. 3***, 2 vs. 4***, 3 vs. 4*** |
| Hippocampal volume, right (mean ml, SD) | 3.97 | 0.43 | 4.20 | 0.37 | 4.43 | 0.36 | 4.06 | 0.36 | 3.74 | 0.33 | *** | 1 vs. 2***, 1 vs. 3***, 1 vs. 4***, 2 vs. 3***, 2 vs. 4***, 3 vs. 4*** |
| Amygdalar volume, left (ml) | 1.68 | 0.19 | 1.77 | 0.18 | 1.88 | 0.14 | 1.71 | 0.17 | 1.58 | 0.15 | *** | 1 vs. 2***, 1 vs. 3*, 1 vs. 4***, 2 vs. 3***, 2 vs. 4***, 3 vs. 4*** |
| Amygdalar volume, right (ml) | 1.50 | 0.16 | 1.58 | 0.16 | 1.67 | 0.14 | 1.53 | 0.14 | 1.43 | 0.13 | *** | 1 vs. 2***, 1 vs. 3*, 1 vs. 4***, 2 vs. 3***, 2 vs. 4***, 3 vs. 4*** |

### Data Processing and Statistical Analysis

Details on all analysis methods can be found in Supplementary Methods (data available from bioRxiv (Additional Methods) https://doi.org/10.1101/239160).

#### Blood pressure classification

For statistical analyses, all available BP measurements per participant were averaged to one mean SBP and DBP, respectively. Based on these averages, we categorized BP according to the European guidelines for the management of arterial hypertension^23^: *category 1* (SBP<120 mmHg and DBP<80 mmHg), *category 2* (SBP 120-129 mmHg or DBP 80-84 mmHg), *category 3* (SBP 130-139 mmHg or DBP 85-89 mmHg) and *category 4* (SBP≥140 mmHg or DBP≥90 mmHg).

#### Voxel-based morphometry (VBM): association of regional GMV and BP within each sample

For each of the four samples, 3 Tesla high-resolution T1-weighted 3-D whole-brain images were processed using VBM and the diffeomorphic anatomical registration using exponentiated lie algebra (DARTEL) method^19,20^ within SPM12. Voxel-wise general linear models (GLM) were performed to relate BP and GMV within each sample: we tested for a continuous relationship between GMV and SBP or DBP, in separate models, with a multiple linear regression *t*-contrast. The overall effect of BP category on GMV was tested with an Analysis of Variance (ANOVA) *F*-contrast. To assess differences in GMV between BP categories, the following pairwise *t*-comparisons were tested: (a) category 4 vs. category 1, (b) category 3 vs. category 1, (c) category 2 vs. category 1. All analyses included total intracranial volume (TIV), sex and age as covariates. The influence of body mass index (BMI) did not significantly contribute to the models and was thus not included as covariate in the analyses. We considered a sample eligible for image-based meta-analysis if its *F*- contrast effects exceeded an uncorrected peak-level threshold of *p*<0.001. Effects within each sample were explored at cluster-level *p*<0.05 with family-wise error correction for multiple comparisons.

#### Image-based meta-analysis (IBMA): association of regional GMV and BP across samples

To evaluate cumulative results from all samples while considering their heterogeneities, we combined the VBM outcome of each sample in IBMA. Meta-analyses were performed on the unthresholded *t*-maps with SDM software using default parameters^24^. Statistical significance of mean effect size maps was evaluated according to validated thresholds of high meta-analytic sensitivity and specificity^24^: voxel threshold=*p*<0.005, peak height threshold=SDM-*Z*>1.0 and cluster extent threshold=*k*≥10 voxels.

Exploratory IBMA for positive associations were performed in analogy to negative associations as described above.

#### IBMA of regions of interest (ROI): association of regional GMV and BP across samples in hippocampus and amygdala

We performed IBMA within atlas-defined masks to test if regional bilateral hippocampal and amygdalar volumes related to SBP, DBP and BP categories, respectively. The statistical thresholds were defined as *p*<0.05, SDM-*Z*>1.0 and *k*≥1 voxels.

#### Volumetry: association of total brain volumes and BP within the pooled sample

In addition to VBM and IBMA, we explored if total brain volumes (average volume over all voxels within a region) differed between BP categories. Specifically, we tested if estimated total intracranial volume, total grey matter volume, total white matter volume (WMV), total amount of white matter hyperintensities (WMH), total cerebrospinal fluid volume (CSFV), total left and right hippocampal and amygdalar volume differed between BP categories. WMH was assessed by Fazekas scale ratings from Fluid-attenuated inversion recovery (FLAIR) images^25^. For these comparisons within the total sample, we defined correlation models (for SBP and DBP as independent variable, respectively) and ANOVA models for BP category as independent variable. The models included the respective volume as dependent variable, as well as TIV (where applicable), sex, age, and sample (where applicable) as covariates. We considered *p*-values<0.05 as significant. The analyses were performed with R (3.2.3, R Core Team, 2015, Vienna, Austria; https://www.R-project.org/).

### Data Sharing

Results (i.e. unthresholded whole-brain statistical maps) from VBM analyses of each sample and from all IBMAs can be found online in the public repository NeuroVault for detailed, interactive inspection (http://neurovault.org/collections/FDWHFSYZ/). Raw data of samples 1-3 available from https://www.openfmri.org/dataset/ds000221/.

## Results

### Sample characteristics

The characteristics of the total sample by BP category are reported in Table 1. The total sample included 423 participants between 19-40 years of whom 177 were women (42%). Mean (SD) age was 27.7 (5.3) years. SBP, DBP, and BMI differed between BP categories (all *p*<0.001). An effect of sex yielded that men were more frequent in higher BP categories (all *p*<0.001).

Table 2 shows differences in characteristics between the four included samples. The samples differed in almost all characteristic variables, specifically regarding sex, age, SBP, DBP, smoking status, and brain volumes (all *p*<0.001).

### VBM: association of regional GMV and BP within each sample

Figure 2 shows differences in regional GMV between BP categories for each of the four samples tested with an ANOVA *F*-contrast. Results show significant clusters of various extents that were distributed heterogeneously between the samples. Exploration of sample-specific effects showed a cluster in the left posterior insula for the *F*-contrast (peak MNI coordinates [−38,−24,24], *F*=11.35, cluster size k=1239) as well as clusters in left inferior frontal gyrus ([−42,34,0], *T*=4.99, k=2039) and in right anterior cingulate cortex ([14,34,14], *T*=4.44, k=2086) for the contrast BP category 4<1 in sample 1. In sample 2, the contrast BP category 4<1 yielded a cluster in left planum polare ([−44,−22,−3], *T*=10.70, k=1151) and the contrast BP category 2<1 yielded a trend for a cluster in left middle temporal gyrus (*p*_*FWE*_=0.059, [−54,−28,−9], *T*=5.41, k=683). Furthermore, higher DBP was associated with a cluster of lower GMV in left middle temporal gyrus in sample 2 ([−57,−45,6], *T*=6.03, k=1180). All other comparisons yielded no suprathreshold voxels (all *p*_*FWE*_>0.05). The statistical maps for sample-specific effects can be inspected on NeuroVault.

**Figure 2.**
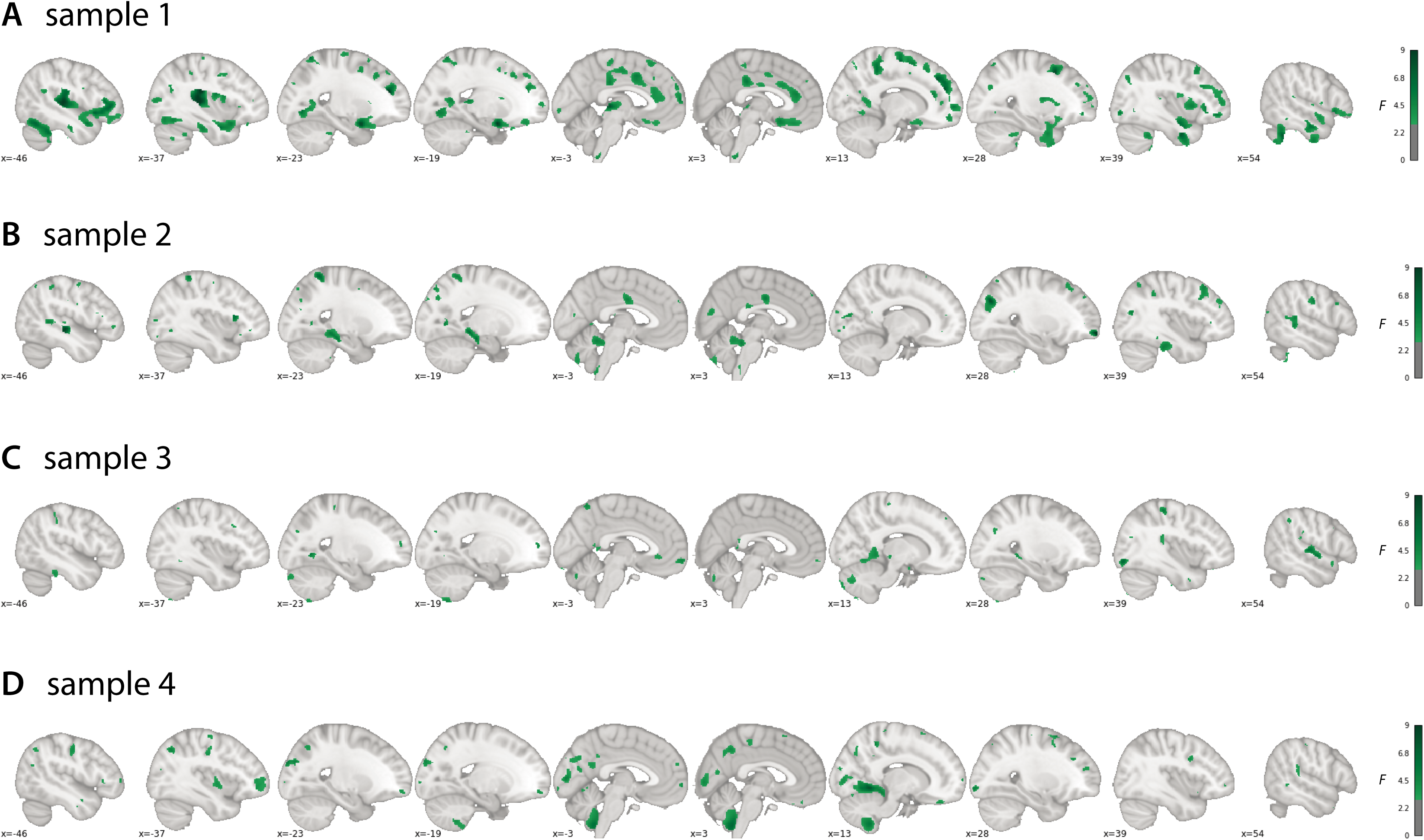
Associations between grey matter volume and blood pressure within each sample. Sagittal views of VBM *F*-contrast results showing the overall effect of BP category on GM volume per sample. Each sample is represented in one row (**A-D**). Slice order runs from left hemisphere (left-hand side of the plot) to right hemisphere (right-hand side of the plot). Color bars represent *F*-values (uncorrected). Sample sizes: sample 1 *n* = 81; sample 2 *n* = 52; sample 3 *n* = 70; sample 4 *n* = 220. 3D-volumetric results of these analyses can be inspected in detail on http://neurovault.org/collections/FDWHFSYZ/. VBM: Voxel-based morphometry. BP: Blood Pressure. GM: Grey Matter.

### IBMA: association of regional GMV and BP across samples

#### Meta-analytic parametric relations between lower GMV and higher BP

As expected, increases in systolic and diastolic BP were associated with lower GMV. Specifically, higher SBP related to lower GMV in right paracentral/cingulate areas ([8,−30,56], SDM-*Z*=−3.859, *k*=288), bilateral inferior frontal gyrus (IFG, left: [−40,30,0], SDM-*Z*=−3.590, *k*=49; right: [−40,30,0], SDM-*Z*=−3.394, *k*=16), bilateral sensorimotor cortex (left: [−58,−20,24], SDM-*Z*=−3.290, *k*=146; right: [48,0,48], SDM-*Z*=−3.196, *k*=127), bilateral superior temporal gyrus (left: [−52,−10,6], SDM-*Z*=−3.268, *k*=78; right: [64,−42,12], SDM-*Z*=−3.192, *k*=42), bilateral cuneus cortex (left: [−8,−76,18], SDM-*Z*=−3.019, *k*=27; right: [10,−68,26], SDM-*Z*=− 2.937, *k*=18), and right thalamus ([8,−28,2], SDM-*Z*=−2.977, *k*=45; Figure 3A, Table 3). Increases in diastolic BP were related to lower GMV in bilateral anterior insula (left: [− 36,26,6], SDM-*Z*=−3.876, *k*=266; right: [34,10,8], SDM-*Z*=−3.139, *k*=100), frontal regions ([−26,24,54], SDM-*Z*=−3.820, *k*=62), right midcingulate cortex ([4,−34,50], SDM-*Z*=−3.545, *k*=246), bilateral inferior parietal areas (left: [−46,−26,48], SDM-*Z*=−3.239, *k*=59; right: [44,- 44,50], SDM-*Z*=−3.188, *k*=18), and right superior temporal gyrus ([60,2,−12], SDM-*Z*=−2.991, *k*=35; Figure 3B, Table 3).

**Table 3.** Image-based meta-analysis results of regional grey matter volume differences associated with blood pressure. Image-based meta-analysis results of significant clusters yielding lower grey matter volume for the respective contrast of interest. Columns indicate cluster-specific MNI coordinates of peak voxels, meta-analytic SDM-*Z*-value, meta-analytic *p*-value, number of voxels in cluster and anatomical label of the peak voxel. Anatomical labels were assigned using SPM’s Anatomy toolbox. *Q* and *I*^*2*^ are measures of meta-analytic heterogeneity. Voxel threshold was set to *p*<0.005, peak height threshold was set to SDM-*Z*>1.0 and cluster extent threshold was set to k≥10 voxels as recommended by ^24^. Final voxel size was 2 × 2 × 2 mm^3^. MNI: Montreal Neurological Institute. SDM: Seed-based *d* Mapping. SBP: Systolic blood pressure. DBP: Diastolic blood pressure.

|  | MNI<br>(x/y/z) | SDM-Z | *P* | *k* | Peak Description | *Q* | *I*<sup>2</sup> |
| --- | --- | --- | --- | --- | --- | --- | --- |
| *Negative Correlation with Systolic Blood Pressure* |  |  |  |  |  |  |  |
|  | 8,-30,56 | -3.859 | 0.000 | 288 | Right paracentral lobule | 0.000 | 0.0 |
|  | -40,30,0 | -3.590 | 0.000 | 49 | Left inferior frontal gyrus (p. triangularis) | 0.053 | 0.0 |
|  | 36,6,34 | -3.394 | 0.000 | 16 | Right inferior frontal gyrus (p. opercularis) | 0.000 | 0.0 |
|  | 10,2,40 | -3.325 | 0.001 | 45 | Right midcingulate cortex | 0.000 | 0.0 |
|  | -58,-20,24 | -3.290 | 0.001 | 146 | Left postcentral gyrus | 0.000 | 0.0 |
|  | -52,-10,6 | -3.268 | 0.001 | 78 | Left superior temporal gyrus | 0.000 | 0.0 |
|  | 48,32,10 | -3.204 | 0.001 | 27 | Right inferior frontal gyrus (p. triangularis) | 0.000 | 0.0 |
|  | 48,0,48 | -3.196 | 0.001 | 127 | Right precentral gyrus | 0.000 | 0.0 |
|  | 64,-42,12 | -3.192 | 0.001 | 42 | Right superior temporal gyrus | 0.000 | 0.0 |
|  | 6,8,-18 | -3.110 | 0.001 | 40 | Right subgenual cingulate cortex | 0.000 | 0.0 |
|  | 50,8,28 | -3.045 | 0.002 | 26 | Right inferior frontal gyrus (p. opercularis) | 0.000 | 0.0 |
|  | -8,-76,18 | -3.019 | 0.002 | 27 | Left cuneus cortex | 0.175 | 0.0 |
|  | 8,-28,2 | -2.977 | 0.002 | 45 | Right thalamus | 0.000 | 0.0 |
|  | 10,-68,26 | -2.937 | 0.002 | 18 | Right cuneus cortex | 0.000 | 0.0 |
|  | 58,4,-8 | -2.934 | 0.002 | 32 | Right temporal pole | 0.000 | 0.0 |
|  | -28,10,60 | -2.896 | 0.003 | 19 | Left middle frontal gyrus | 0.000 | 0.0 |
|  | -52,-12,42 | -2.860 | 0.003 | 10 | Left postcentral gyrus | 0.116 | 0.0 |
| *Negative Correlation with Diastolic Blood Pressure* |  |  |  |  |  |  |  |
|  | -36,26,6 | -3.876 | 0.000 | 266 | Left insula | 0.000 | 0.0 |
|  | -26,24,54 | -3.820 | 0.000 | 62 | Left middle frontal gyrus | 0.000 | 0.0 |
|  | 4,-34,50 | -3.545 | 0.000 | 246 | Right midcingulate cortex | 0.000 | 0.0 |
|  | -60,-24,14 | -3.462 | 0.000 | 90 | Left supramarginal gyrus | 0.000 | 0.0 |
|  | -46,-26,48 | -3.239 | 0.001 | 59 | Left inferior parietal lobule | 0.000 | 0.0 |
|  | 44,-44,50 | -3.188 | 0.001 | 18 | Right inferior parietal lobule | 0.257 | 0.0 |
|  | 36,8,32 | -3.180 | 0.001 | 25 | Right inferior frontal gyrus (p. opercularis) | 0.000 | 0.0 |
|  | 34,10,8 | -3.139 | 0.001 | 100 | Right insula | 0.000 | 0.0 |
|  | 28,14,60 | -3.069 | 0.001 | 12 | Right superior frontal gyrus | 0.000 | 0.0 |
|  | 62,-44,16 | -2.991 | 0.002 | 35 | Right superior temporal gyrus | 0.000 | 0.0 |
|  | -38,14,-20 | -2.983 | 0.002 | 30 | Left temporal pole | 0.000 | 0.0 |
|  | 60,2,-12 | -2.862 | 0.003 | 13 | Right superior temporal gyrus | 0.189 | 0.0 |
|  | -38,40,32 | -2.845 | 0.003 | 14 | Left middle frontal gyrus | 0.000 | 0.0 |
|  | 30,28,0 | -2.796 | 0.003 | 14 | Right insula | 0.000 | 0.0 |
|  | -36,8,10 | -2.788 | 0.003 | 24 | Left insula | 0.000 | 0.0 |
|  | -58,-46,30 | -2.750 | 0.004 | 11 | Left supramarginal gyrus | 0.000 | 0.0 |
|  | -34,32,32 | -2.734 | 0.004 | 11 | Left middle frontal gyrus | 0.201 | 0.0 |
| *Category 4 (SBP≥140 mmHg or DBP≥90 mmHg) < Category 1 (SBP<120 mmHg and DBP<80 mmHg)* |  |  |  |  |  |  |  |
|  | -52,28,12 | -3.473 | 0.000 | 107 | Left inferior frontal gyrus (p. triangularis) | 0.324 | 3.9 |
|  | -48,-4,4 | -3.322 | 0.000 | 93 | Left rolandic operculum | 0.297 | 1.8 |
|  | 18,-52,-48 | -3.097 | 0.001 | 40 | Right cerebellum, hemispheric lobule VIIIb | 0.000 | 0.0 |
|  | 40,30,26 | -3.093 | 0.001 | 10 | Right inferior frontal gyrus (p. triangularis) | 0.000 | 0.0 |
|  | 48,32,10 | -3.064 | 0.001 | 48 | Right inferior frontal gyrus (p. triangularis) | 0.000 | 0.0 |
|  | -38,48,-16 | -3.014 | 0.001 | 40 | Left inferior frontal gyrus (p. orbitalis) | 0.000 | 0.0 |
|  | -54,-12,42 | -2.940 | 0.002 | 30 | Left postcentral gyrus | 0.000 | 0.0 |
|  | -8,-76,18 | -2.936 | 0.002 | 14 | Left cuneus | 0.000 | 0.0 |
|  | -16,-36,-18 | -2.872 | 0.002 | 24 | Left cerebellum, hemispheric lobule V | 0.000 | 0.0 |
|  | 12,-42,48 | -2.854 | 0.002 | 11 | Right midcingulate cortex | 0.000 | 0.0 |
|  | -12,-50,-56 | -2.849 | 0.002 | 30 | Left cerebellum, hemispheric lobule IX | 0.325 | 4.0 |
|  | 10,-52,18 | -2.836 | 0.002 | 21 | Right precuneus | 0.000 | 0.0 |
|  | 64,-44,14 | -2.824 | 0.002 | 26 | Right superior temporal gyrus | 0.000 | 0.0 |
|  | 10,-66,28 | -2.821 | 0.002 | 56 | Right precuneus | 0.000 | 0.0 |
|  | 6,-28,50 | -2.792 | 0.003 | 16 | Right midcingulate cortex | 0.025 | 0.0 |
|  | 18,-54,22 | -2.765 | 0.003 | 15 | Right precuneus | 0.000 | 0.0 |
| Category 3 (SBP 130-139 mmHg or DBP 85-89 mmHg) < Category 1 | 36,6,34 | -3.474 | 0.000 | 179 | Right inferior frontal gyrus (p. opercularis) | 0.000 | 0.0 |
|  | 6,-28,54 | -3.119 | 0.000 | 179 | Right posterior-medial frontal gyrus | 0.127 | 0.0 |
|  | 48,-50,20 | -2.917 | 0.000 | 74 | Right middle temporal gyrus | 0.000 | 0.0 |
|  | -60,-20,36 | -2.857 | 0.000 | 205 | Left postcentral gyrus | 0.000 | 0.0 |
|  | -40,30,2 | -2.598 | 0.000 | 24 | Left inferior frontal gyrus (p. triangularis) | 0.000 | 0.0 |
|  | 36,8,-18 | -2.523 | 0.001 | 123 | N/A (Right insula) | 0.000 | 0.0 |
|  | 42,-74,12 | -2.454 | 0.001 | 25 | Right middle occipital gyrus | 0.200 | 0.0 |
|  | -62,-42,28 | -2.433 | 0.001 | 41 | Left supramarginal gyrus | 0.000 | 0.0 |
|  | 20,-32,6 | -2.384 | 0.001 | 133 | Right thalamus | 0.000 | 0.0 |
|  | -10,36,-6 | -2.384 | 0.001 | 102 | Left anterior cingulate cortex | 0.000 | 0.0 |
|  | 28,-94,-4 | -2.373 | 0.001 | 14 | Right inferior occipital gyrus | 0.000 | 0.0 |
|  | -12,-32,0 | -2.264 | 0.002 | 133 | Left thalamus | 0.237 | 0.0 |
|  | -56,-64,16 | -2.222 | 0.002 | 28 | Left middle temporal gyrus | 0.050 | 0.0 |
|  | -40,8,30 | -2.197 | 0.002 | 82 | Left precentral gyrus | 0.000 | 0.0 |
|  | -12,-54,14 | -2.187 | 0.002 | 20 | Left precuneus | 0.000 | 0.0 |
| Category 2 (SBP 120-129 mmHg or DBP 80-84 mmHg) < Category 1 | -54,-10,14 | -3.407 | 0.000 | 230 | Left rolandic operculum | 0.016 | 0.0 |
|  | 30,-96,-8 | -3.290 | 0.000 | 102 | Right inferior occipital gyrus | 0.038 | 0.0 |
|  | -34,-16,-30 | -3.164 | 0.000 | 133 | Left fusiform gyrus | 0.000 | 0.0 |
|  | -8,-54,22 | -3.084 | 0.000 | 433 | Left precuneus | 0.000 | 0.0 |
|  | 54,-24,32 | -2.968 | 0.000 | 31 | Right supramarginal gyrus | 0.019 | 0.0 |
|  | -46,28,0 | -2.942 | 0.000 | 41 | Left inferior frontal gyrus (p. triangularis) | 0.000 | 0.0 |
|  | -64,-20,30 | -2.939 | 0.000 | 227 | Left postcentral gyrus | 0.000 | 0.0 |
|  | -62,-42,34 | -2.876 | 0.000 | 68 | Left supramarginal gyrus | 0.000 | 0.0 |
|  | -36,-64,42 | -2.827 | 0.001 | 30 | Left angular gyrus | 0.000 | 0.0 |
|  | 18,-72,54 | -2.804 | 0.001 | 40 | Right superior parietal lobule | 0.083 | 0.0 |
|  | 46,-74,12 | -2.734 | 0.001 | 26 | Right middle temporal gyrus | 0.000 | 0.0 |
|  | 8,-18,46 | -2.647 | 0.001 | 32 | Right midcingulate cortex | 0.000 | 0.0 |
|  | 56,-32,12 | -2.470 | 0.002 | 52 | Right superior temporal gyrus | 0.000 | 0.0 |
|  | 28,-72,-38 | -2.413 | 0.002 | 23 | Right cerebellum, crus I | 0.000 | 0.0 |
|  | -62,-22,-30 | -2.403 | 0.003 | 11 | N/A (Left inferior temporal gyrus) | 0.000 | 0.0 |

**Figure 3.**
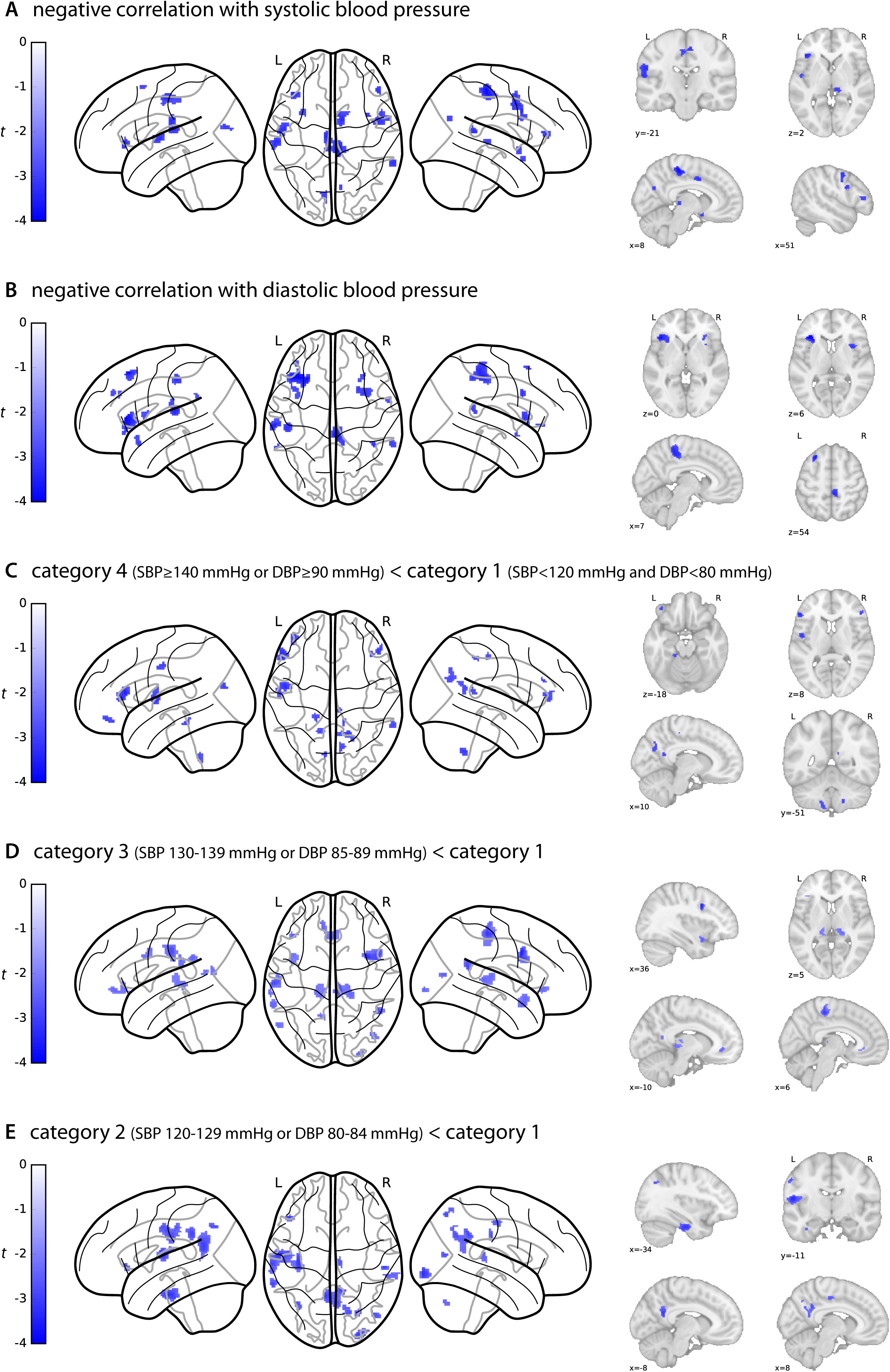
Meta-analytic differences in grey matter volume between blood pressure categories. Glass brain views of image-based meta-analysis results for the blood pressure category contrasts of interest with relevant slice views below (**A-E**). **A** and **B** depict associations between higher SBP/DBP, respectively, and lower gray matter volume, i.e. negative correlations. Blue clusters indicate meta-analytic grey matter volume differences for the given contrast at a voxel threshold of *p*<0.005 with peak height threshold of SDM-*Z*<-1.0 and cluster extent threshold of *k* ≥10 (validated for high meta-analytic sensitivity and specificity^24^). Color bars represent SDM-*Z*-values. 3D-volumetric results of these analyses can be inspected in detail on http://neurovault.org/collections/FDWHFSYZ/. SDM: Seed-based *d* Mapping. SBP: Systolic blood pressure. DBP: Diastolic blood pressure. L: Left hemisphere. R: Right hemisphere.

#### Meta-analytic differences in regional GMV between BP categories

Meta-analytic results for category 4 (highest BP) compared to category 1 (lowest BP) yielded lower regional GMV in frontal, cerebellar, parietal, occipital, and cingulate regions (Figure 3C). Table 3 describes the specific regions with lower GMV, including bilateral IFG (left: [−52,- 28,12], SDM-*Z*=−3.473, *k*=107; right: [40,30,26], SDM-*Z*=−3.093, *k*=10), right midcingulate cortex ([12,−42,48], SDM-*Z*=−2.854, *k*=11), and right precuneus ([10,−52,18], SDM-*Z*=−2.836, *k*=21).

We also compared GMV of individuals at sub-hypertensive levels (category 3 and 2, respectively) to GMV of individuals in category 1. Figure 3D shows meta-analysis results for the comparison between category 3 and category 1. Compared to category 1, category 3 was associated with lower GMV in bilateral IFG (left: [−40,30,2], SDM-*Z*=−2.598, *k*=24; right: [36,6,34], SDM-*Z*=−3.474, *k*=179), sensorimotor cortices (left: [−60,−20,36], SDM-*Z*=−2.857, *k*=205; right: [6,−28,54], SDM-*Z*=−3.119, *k*=179), bilateral middle temporal gyrus (left: [−56,−64,16], SDM-*Z*=−2.222, *k*=28; right: [48,−50,20], SDM-*Z*=−3.119, *k*=179), right insula ([36,8,−18], SDM-*Z*=−2.523, *k*=123), right occipital regions ([42,−74,12], SDM-*Z*=−2.454, *k*=25), left parietal ([−60,−20,36], SDM-*Z*=−2.857, *k*=205), bilateral thalamus (left: [−12,−32,0], SDM-*Z*=− 2.264, *k*=133; right: [20,−32,6], SDM-*Z*=−2.384, *k*=133), left anterior cingulate cortex ([−10,36,- 6], SDM-*Z*=−2.384, *k*=102), and left precuneus ([−12,−54,14], SDM-*Z*=−2.187, *k*=20; Table 3).

Figure 3E illustrates brain regions that yielded lower meta-analytic GMV comparing category 2 to category 1. These include left frontal regions ([−54,−10,14], SDM-*Z*=−3.407, *k*=230), right inferior occipital gyrus ([30,−96,−8], SDM-*Z*=−3.290, *k*=102), bilateral temporal regions (left: [− 34,−16,−30], SDM-*Z*=−3.164, *k*=133; right: [46,−74,12], SDM-*Z*=−2.734, *k*=26), left precuneus ([−8,−54,22], SDM-*Z*=−3.084, *k*=433) and inferior parietal regions (supramarginal, [54,−24,32], SDM-*Z*=−2.968, *k*=31, and angular gyri, [−36,−64,42], SDM-*Z*=−2.827, *k*=30), as well as midcingulate cortex ([8,−18,46], SDM-*Z*=−2.647, *k*=32; Table 3).

#### Meta-analytic differences in regional hippocampal and amygdalar volumes between BP categories

In this IBMA ROI comparison, higher SBP was correlated with lower bilateral posterior medial hippocampal volume (Figure 4). Higher DBP was correlated with lower left hippocampal volume and lower right anterior hippocampal volume. Furthermore, all higher BP categories were associated with lower regional hippocampal volume when compared to the lowest BP category 1 (Figure 4). Compared to category 1, BP category 4 was predominantly associated with lower left medial posterior hippocampus volume and category 3 with lower bilateral posterior and left medial hippocampus volume across samples. Lower volume comparing categories 2 and 1 was predominantly located in left lateral anterior hippocampus. Category 4 vs. category 1 and the associations with higher SBP and DBP also yielded significantly lower regional volume in bilateral amygdala, respectively. Effect sizes highly varied across samples (Figure 4).

**Figure 4.**
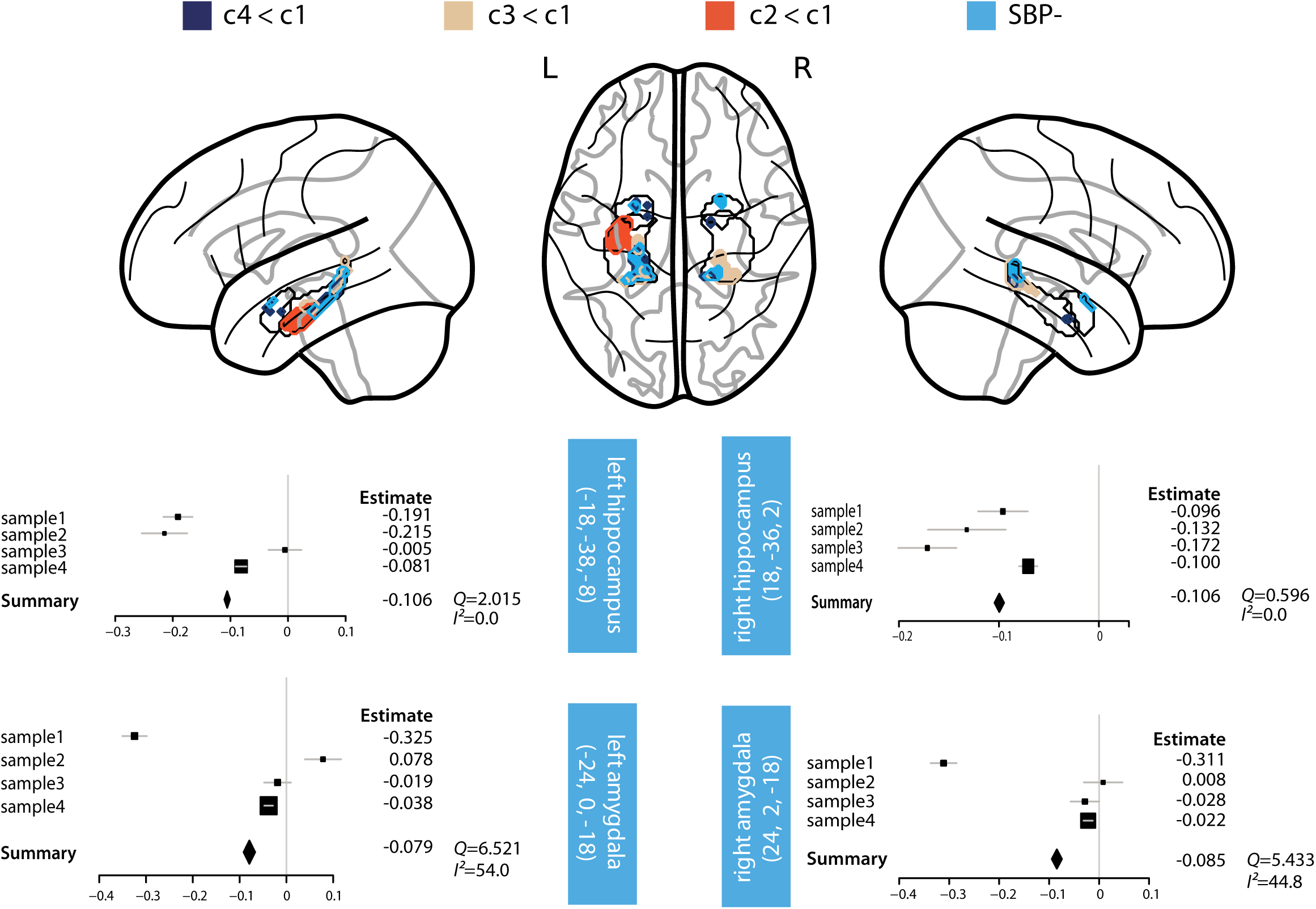
Meta-analytic differences in volumes of hippocampus and amygdala (Region-of-Interest analysis). Upper part of plot: Glass brain views of image-based meta-analysis ROI results for the blood pressure category contrasts of interest in bilateral hippocampus and amygdala masks. Voxel threshold was set to *p*<0.05 with a peak height threshold of SDM-*Z*<-1.0 and a cluster extent threshold of *k* ≥1. Lower part of plot: Exemplary forest plots of sample-specific peak voxels’ effect sizes for the negative correlation with SBP in the respective ROI. The box sizes are determined by each sample’s weight. Light blue boxes include ROI name and MNI coordinates of the peak voxel. *Q* and *I*^*2*^ are measures of meta-analytic heterogeneity. Definition of blood pressure categories: *category 1* (SBP<120 mmHg and DBP<80 mmHg), *category 2* (SBP 120-129 mmHg or DBP 80-84 mmHg), *category 3* (SBP 130-139 mmHg or DBP 85-89 mmHg) and *category 4* (SBP≥140 mmHg or DBP≥90 mmHg). SBP-: negative correlation with SBP. ROI: Region of Interest. SDM: Seed-based *d* Mapping. MNI: Montreal Neurological Institute. SBP: Systolic blood pressure. DBP: Diastolic blood pressure. L: Left hemisphere. R: Right hemisphere.

#### Meta-analytic relations between higher GMV and higher BP

Exploratory analyses also revealed associations between higher BP and higher GMV (data available from bioRxiv (Supplementary Figure 1, Supplementary Table 2) https://doi.org/10.1101/239160, and NeuroVault maps). However, the cumulative positive effects are comparably weaker than the cumulative negative results (Table 3, negative: 17 out of 34 clusters from parametric analyses with SDM-Z>3.0; positive: 0 out of 28 clusters from parametric analyses with SDM-Z>3.0), they show greater heterogeneity across studies (negative: maximum *I*^*2*^=4.0; positive: maximum *I*^*2*^=56.3) and they seem to appear primarily in regions where standard preprocessing of brain tissue is suboptimal (e.g. in cerebellum/inferior occipital regions^26^). We therefore regard these findings as overall questionable. By also providing the results as statistical maps on NeuroVault, future investigations can use the data for reliability analyses of potential positive associations.

### Volumetry in pooled sample: association of total brain volumes and BP

All associations of volumetric brain measures (TIV, total GMV, total, WMV, total CSFV, total hippocampal, total amygdalar volume and total WMH) with SBP or DBP in the correlation models, or with BP categories in the ANOVA models were below the statistical threshold (all *p*>0.05, Table 1).

## Discussion

In this image-based meta-analysis of four previously unpublished independent samples, we found that elevated, sub-hypertensive BP was correlated with lower GMV in several brain regions, including parietal, frontal, and subcortical structures in young adults (<40 years). These regions are consistent with the lower regional GMV observed in middle-aged and older individuals with HTN^5,6,9–11,15^. Our results show that BP-associated GM alterations emerge earlier in adulthood than previously assumed and continuously across the range of BP.

Interestingly, we found that BP was associated with lower hippocampal volume. In older individuals, the hippocampal formation and surrounding structures are known to be affected by HTN^5,8–10,15^. In a meta-analytic evaluation of HTN-effects on total GMV and on hippocampal volume, lower volumes across studies were only consistently found for the hippocampus^15^. In analogy to those findings, our results showed that hippocampal volume was affected by higher BP in a considerably younger sample. It should be mentioned that the effects in hippocampus only exceeded statistical thresholds in ROI analyses, similar to previous reports of lower hippocampal volume in older samples with manifest HTN that were all ROI-based^5,9,10,15^. As potential pathophysiological explanations it has been proposed that medial temporal (and frontal regions) might be especially sensitive to effects of pulsation, hypoperfusion and ischemia, which often result from increasing pressure^3,15^.

We furthermore observed correlations between lower amygdalar and thalamic volumes and higher BP, notably already below levels which are currently regarded as hypertensive. Amygdalar and thalamic nuclei are substantially involved in BP regulation as they receive baroreceptor afferent signals via the brainstem and mesencephalic nuclei, relaying these signals to primary cortical regions of viscerosensory integration, such as anterior cingulate cortex and insula^27^. It has been shown that lower amygdalar volume correlates with increased BP-reactivity during cognitive demand among young normotensive adults^18^. Previous studies have reported lower thalamic volume in HTN^5^, heart failure^28^, asymptomatic carotid stenosis^29^, and aging^30^. Higher systolic BP has also been related to higher mean diffusivity of white matter thalamic radiations^13^. Our results are in line with accumulating evidence of amygdalar and thalamic involvement in cardiovascular (dys-) regulation but may also reflect early pathology in these regions. For example, occurrence of neurofibrillary tangles in thalamus has also been reported in the earliest stages of AD neuropathology^34^.

Beyond subcortical structures, we found lower volumes in cortical regions: cingulate volume and insular volume were markedly lower with higher DBP in the meta-analysis results and in the individual analyses of sample 1. As noted above, these regions constitute primary cortical sites of afferent viscerosensory integration and modulate homeostasis via efferents to brainstem nuclei^27^. Lesions in cingulate cortex and insula result in altered cardiovascular regulation, increased sympathetic tone^31,32^ and myocardial injury^33^. Both regions are also critical for the appraisal and regulation of emotion and stress^27^. Thus, structural alterations in these regions may contribute to insidious BP elevations via sympathetic pathways.

Frontal and parietal volumes were affected in all our statistical comparisons. The precuneus cortex, especially, was associated with lower GMV in BP categories 4, 3, and 2 compared to category 1. Our results of lower BP-related GMV in regions such as hippocampal, frontal and parietal areas highlight specific brain regions which are known to be vulnerable to putative vascular or neurodegenerative damage mechanisms^5,6,8–11,15,35^. Raised midlife BP is not only known to be a major risk factor for vascular dementia, but some reports suggest a link between HTN and AD-type pathophysiology^3,4^. For example in neuropathological studies, raised midlife BP has been associated with lower post-mortem brain weight, increased numbers of hippocampal neurofibrillary tangles, and higher numbers of hippocampal and cortical neuritic plaques^8^. Similarly, a potential pathophysiological link between HTN and AD has been supported by noninvasive MRI studies: regions referred to as AD-signature regions (including inferior parietal, precuneus cortices, and medial temporal structures) have been associated with cortical thinning years before clinical AD-symptoms arise^35^ and with brain volume reductions predicted by increasing BP from middle to older age^5^. In light of these previous results, our findings of lower BP-related GMV in AD-signature regions may be indicative of a link to AD-pathology at an even earlier age; however, this cannot be causally inferred from our cross-sectional data. In the study by Power et al.^5^, BP also predicted volume loss in non-AD-typical brain regions, such as frontal lobe and subcortical gray matter, which may relate to other (than AD-related) pathophysiological mechanisms. A similar pattern seems to be reflected in our findings of lower GMV related to higher BP in non-AD-typical regions.

Some previous studies did not find relations between HTN and lower brain volumes, but associated HTN with other forms of structural or functional brain alterations, such as white matter injury^36^ or reduced cerebral perfusion^37^. A key aspect of diverging results is the heterogeneity of methods used to assess brain volumes. Earlier investigations of BP effects on brain tissues have applied manual or automated volumetric methods to quantify total brain volumes in pre-selected ROIs^9,10,12^. The focus of this study was to employ computational anatomy methods to assess *regional GM differences across the whole brain*. We found significant differences between BP groups using VBM but not in the analysis of total brain volumes. This supports the view that VBM is a sensitive measure to quantify regional morphological differences^38^ which might be undetected from the analysis of total brain volumes alone. In addition, we employed random-effects IBMA which results in effects that are consistent across studies and that may otherwise be neglected at sub-threshold. Investigating effects of BP on regional vs. total brain volumes at all stages of health and disease thus warrants further research with standardized methods to identify neuropathological mechanisms.

Our data, however, do not allow inference on causality between lower brain volumes and HTN, which likely involves complex interactions of different pathophysiological mechanisms that still need to be fully elucidated. It is assumed that vascular stiffness, endothelial failure and a dysfunctional blood-brain barrier are precursors of cerebral small and large vessel disease that reduce cerebral blood flow, disturb autoregulatory adjustment and decrease vasomotor reactivity, which may impair perivascular central nervous waste clearance systems^3^. These mechanisms have also been suggested to potentially underlie the epidemiological connection between vascular risk factors, such as HTN, and AD^3^. The similarities between our findings and AD-signature regions (see above) would also be consistent with this putative link. Consequently, demyelination, apoptosis and intoxication of neurons and glial cells, as well as grey and white matter necrosis accumulate and may be reflected in neuroimaging on a macroscopic scale. Lower GMV assessed by VBM, as reported in our study, can thus arise from neuronal loss, but also from alterations of glial cells or composition of microstructural or metabolic tissue properties ^39^. Our findings point to an early effect of such mechanisms on GM integrity which is present in the absence of overt disease, such as HTN, and in young age. Indicators of early atherosclerosis in major peripheral arteries can already be detected in youth^40^. Recently, arterial stiffness has also been associated with WM and GM alterations among adults between 24 and 76 years of age^41^. Thus, already early and subtle vascular changes, deficient cerebral perfusion and impaired perivascular clearance systems may initiate and sustain neuropathology from early to late adulthood.

The cross-sectional design of our four study samples limits the interpretation frame for the results presented. Causality between BP and potential brain damage cannot be assessed with these data but is crucial for implications of early signs of cerebrovascular disease. Furthermore, the study samples differed regarding recruitment, sex distribution, sample size, prevalence of high BP, and data acquisition methods (BP and MRI) which might not represent the general population or standard acquisition protocols: similar to German prevalence^42^, men had higher BP in our study. We thus included sex as covariate in all our analyses to adjust for sex effects. We did not perform separate analyses for men and women given that one of the four samples included only men. However, the topic of sex differences in brain structure related to BP is a very interesting open question for future investigations. In sample 3, only one BP measurement was recorded which could be biased due to white coat hypertension or BP variability. Practice guidelines recommend an average of ≥2 seated readings obtained on ≥2 occasions to provide a more accurate estimate of an individual’s BP level^23,43,44^. By combining the samples in random-effects IBMA, we considered the limitations of each sample and accounted for within- and between-sample heterogeneity and evaluated effects cumulatively. Moreover, this approach enabled us to investigate the expected small effects of BP-related GM alterations in a well-powered total sample of over 400 young adults. To further ensure that the results are not substantially influenced by the heterogeneity of BP measurements across studies, we recalculated the parametric SBP analysis (Figure 3A) with only the first SBP reading in each study. The results of this additional analysis are strikingly similar to the results reported here (data available from bioRxiv (Supplementary Table 3, Supplementary Figure 2) https://doi.org/10.1101/239160). HTN is also the most important risk factor for WM damage^3,12^ and sub-clinical WM injury in relation to elevated BP levels has recently been reported in 19- to 63-year-old adults^13^. As our study included only GM measures, we cannot assess mediating effects of WM injury on GMV differences. We did not observe any significant differences in Fazekas scores for WMH between BP categories, likely due to their lower sensitivity and poorer specificity as a proxy for vascular disease in a sub-clinical sample of young adults with (mostly) normal BP.

Our study shows that BP-related brain alterations may occur in early adulthood and at BP levels below current thresholds for manifest HTN. Contrary to assumptions that BP-related brain damage arises over years of manifest disease our data suggest that subtle pressure-related GM alterations can be observed in young adults without previously diagnosed HTN. Considering our results, large-scale cohort studies should investigate whether sub-hypertensive BP and related brain changes in early adulthood increase the risk for subsequent development of CVD later in life. Gaining insights whether and how the brain is globally affected by vascular changes or if these are specific to susceptible regions could help identifying neuroimaging biomarkers for the earliest stages of CVD. Such data would provide evidence for future guidelines to formulate informed recommendations for BP-management in young adults, which are critical for the prevention of CVD. Lifestyle interventions and neurobehavioral therapy have recently been suggested to benefit CVD prevention^17^. Our results highlight the importance of taking BP levels as a continuous measure into consideration which could help initiate such early preventive measures.

## Supplementary Data

Additional Methods

Supplementary analysis of first blood pressure reading

References

Supplementary Table 1: List of exclusion criteria for each study from which the samples were drawn

Supplementary Table 2: Positive image-based meta-analysis results of regional grey matter volume differences associated with blood pressure.

Supplementary Table 3 – Image-based meta-analysis results of regional grey matter volume differences associated with first systolic blood pressure.

Supplementary Figure 1: Meta-analytic positive differences in grey matter volume between blood pressure categories.

Supplementary Figure 2 – Associations between grey matter volume and first systolic BP reading

### Methods

#### Additional preprocessing steps for MP2RAGE images

Before segmentation, T1-weighted images acquired with an MP2RAGE sequence were additionally masked to remove noise signal outside of the brain (following the procedure described in ^2^): a binarized brain mask was created from the second inversion-contrast volume by setting voxels with intensities of less than 10% of the maximum signal to zero. Any holes in the mask were filled. For the final image, the mask was multiplied with the T1-weighted volume which eliminated background noise but preserved signals from the brain and other tissues. All of these steps were performed with tools in FSL 5.0^3^ (www.fmrib.ox.ac.uk/fsl).

T1-weighted images from MP2RAGE are free of magnetic field inhomogeneity (they are also named *uniform*). Thus, a correction for this bias was omitted and only applied to MPRAGE images. Bias correction for MPRAGE images followed the default settings within SPM12’s *segment* batch. All other processing steps were identical for the pulse sequences.

#### Voxel-based morphometry (VBM) and statistical analysis of regional GMV and BP within each sample

T1-weighted images were processed by using voxel-based morphometry (VBM) and the diffeomorphic anatomical registration using exponentiated lie algebra (DARTEL) method ^4,5^ within SPM12 (12.6685, Wellcome Trust Centre for Neuroimaging, UCL, London, UK; http://www.fil.ion.ucl.ac.uk/spm12/) running under Matlab 9.0.0 (R2016a, MathWorks, Natick, MA, USA). In short, processing of grey matter volume (GMV) probabilities included segmentation into tissue types, sample-specific DARTEL template creation, modulation of grey matter voxels to preserve tissue properties, normalization to MNI space, and 8 mm full-width-at-half-maximum Gaussian smoothing.

Voxel-wise statistical tests were performed in SPM12 to relate BP and GMV within each sample. We first performed a whole-brain analysis, testing for a continuous relationship between GMV and SBP and DBP, respectively, with a linear regression t-contrast. Next we tested for differences in GMV between BP categories. The general linear model for this whole-brain analysis included a factor for BP as variable of interest (levels: (1) category 4, (2) category 3, (3) category 2, (4) category 1). Within each sample, the overall effect of BP category on GMV was tested with an Analysis of Variance (ANOVA) *F*-contrast and the following *t*-contrasts tested pairwise comparisons of interest: (1) category 4 vs. category 1, (2) category 3 vs. category 1, (3) category 2 vs. category 1. All models included total intracranial volume (TIV), sex and age as covariates of no interest. Each *t*- contrast was tested in negative and positive direction (i.e. category A<category B and category A>category B). Data on all effects can be found online in the public repository NeuroVault^6^ (http://neurovault.org/collections/FDWHFSYZ/).

#### Image-based meta-analysis (IBMA): association of regional GMV and BP across samples

To evaluate cumulative results from all samples, we combined the results of each sample in an image-based meta-analysis. The meta-analysis was performed with Anisotropic Effect-Size Signed Differential Mapping (AES-SDM) implemented in the SDM software package using default parameters^7^ (http://www.sdmproject.com/, http://www.sdmproject.com/software/tutorial.pdf). For each sample and *t*- contrast, unthresholded whole-brain *t*-statistic maps were converted to Hedges’ *g* effect size maps and variance maps. To assess weighted mean differences in grey matter across all samples, a meta-analytic model was set up for each voxel. Within this random-effects model, samples are weighted by their sample size, within-study variance and between-study heterogeneity. The result is a mean map of *z*-values which are quotients of the mean effect-sizes and their standard errors. Since these *z*-values are not normally distributed, null distributions were estimated empirically by Monte Carlo randomizations. Voxels in the mean map were randomly permuted within a software-implemented grey matter mask to create null distributions for the assessment of critical *z*-values. We applied 50 permutations, while statistical stability has been shown from 20 permutations on ^7^. Statistical significance was evaluated according to validated thresholds of high meta-analytic sensitivity and specificity^7^: voxel threshold=*p*<0.005, peak height threshold=SDM-*Z*>1.0 and cluster extent threshold=*k* ≥10 voxels. Anatomy toolbox^8^ (version 2.2 for SPM8) was used to automatically label significant clusters in all analyses. Nilearn^9^ (version 0.2.6, https://nilearn.github.io/index.html) was used to visualize statisticalbrain maps.

#### IBMA of regions of interest (ROI): association of regional GMV and BP across samples in hippocampus and amygdala

With the meta-analysis approach, we also tested if specific regions of interest (ROI) that included bilateral hippocampus and bilateral amygdala differed in their volumes related to SBP, DBP and between BP categories. Separate IBMAs were calculated within binary atlas-defined masks for bilateral hippocampus and bilateral amygdala that were retrieved from the latest available version of the Anatomy toolbox^8^ (2.2 for SPM8). Peak voxels’ effect sizes were extracted with SDM software’s *Extract* option and plotted as forest plots (Figure 4) with R (3.2.3, R Core Team, 2015, Vienna, Austria; https://www.R-project.org/) and the package *rmeta* (2.16).

### Supplementary analysis of first blood pressure reading

As correctly pointed out by a reviewer, intra-individual blood pressure variation and white coat hypertension are important to consider when measuring blood pressure^10^. For the meta-analyses, we had reasoned that the most accurate and generalizable basis for estimation of individual BP is the inclusion of all available BP readings within each sample for each participant (i.e. following the intention of the BP measurement in each included study). Since BP measurement protocols differed between the study samples, this approach resulted in the inclusion of BP values from different numbers of BP measurements (see Methods for details): in samples 1, 2 and 4, ≥2 BP readings were taken in varying time intervals and averaged for analyses (n=353). In sample 3, only one BP reading was available and used for analyses (n=70). We accounted for sample differences by employing random-effects models in image-based meta-analyses^7,11^.

Alternatively, by analyzing the first BP reading only, an important difference between the studies would be eliminated, while a potential white coat effect would be possibly emphasized. Following the reviewer’s suggestion, we recalculated one crucial parametric analysis with only the first systolic BP reading in each study. We chose to reanalyze this contrast since it had the largest difference between the first and average BP readings (please note that even in this case the readings overall correlated strongly, Supplementary Figure 2C).

The results of this analysis and the results of the previous analysis (with averaged SBP measurements, as reported in the main manuscript text) are presented in Supplementary Table 3 and Supplementary Figure 2A-B for comparison. The results are strikingly similar: for the analysis based on just the first BP reading, several clusters even showed stronger statisticalresults.

## Supplementary Tables

**Supplementary Table (eTable) 1.**
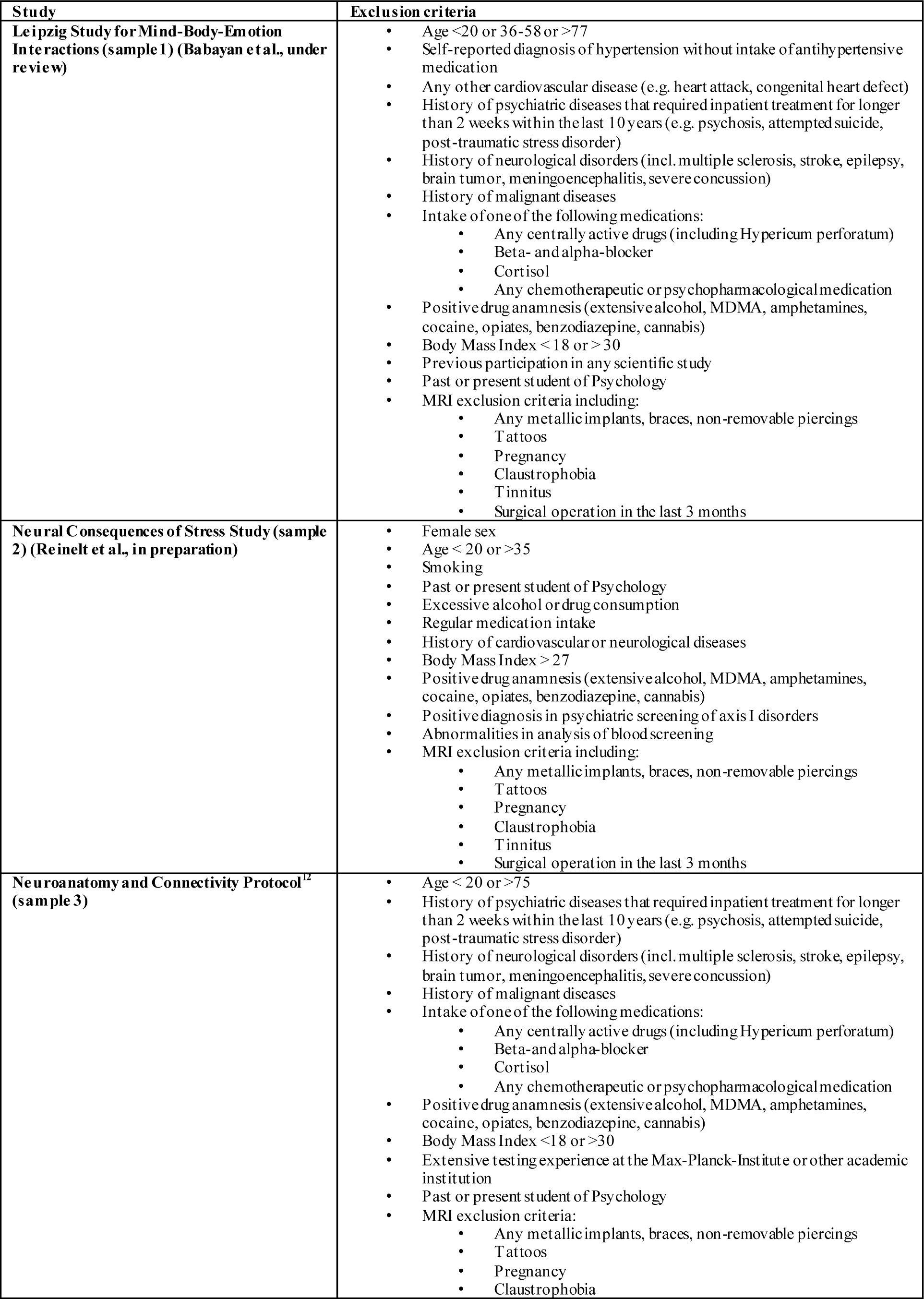
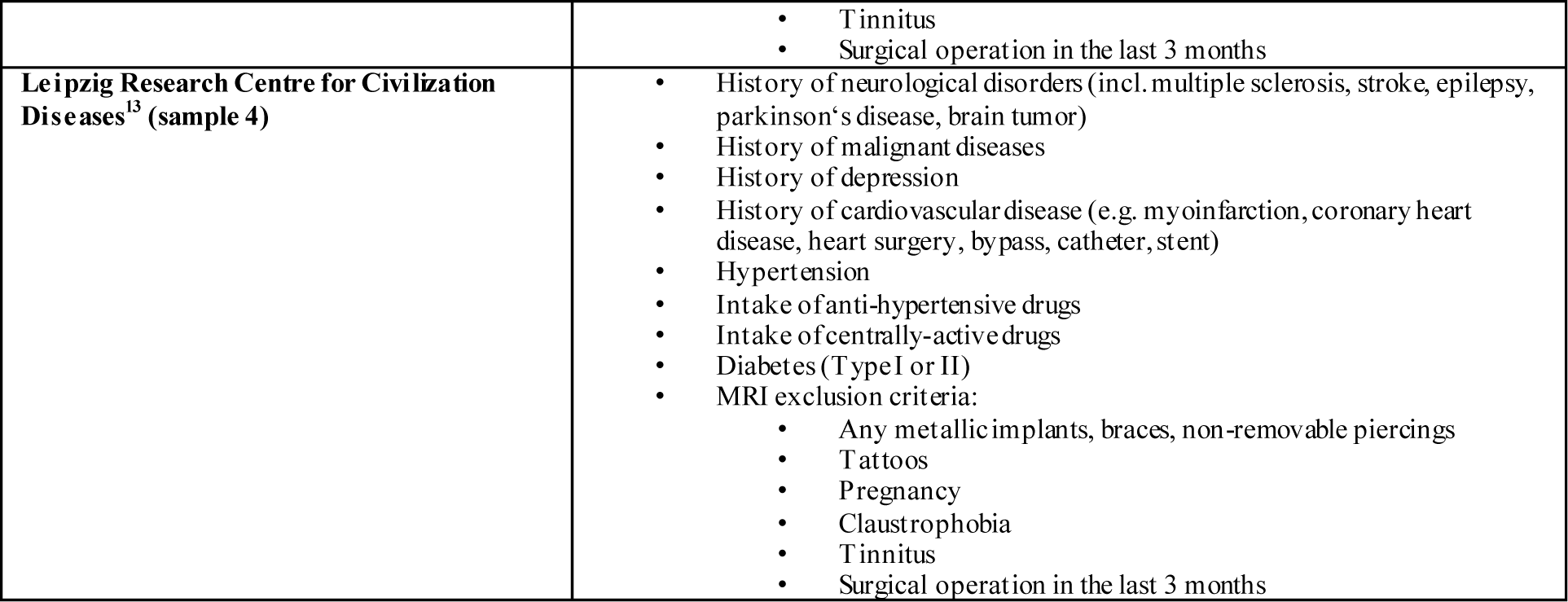
List of exclusion criteria for each study from which the samples were drawn.

| Study | Exclusion criteria |
| --- | --- |
| **Leipzig Study for Mind-Body-Emotion Interactions (sample 1) (Babayan et al., under review)** | • Age <20 or 36-58 or >77<br>• Self-reported diagnosis of hypertension without intake of antihypertensive medication<br>• Any other cardiovascular disease (e.g. heart attack, congenital heart defect)<br>• History of psychiatric diseases that required inpatient treatment for longer than 2 weeks within the last 10 years (e.g. psychosis, attempted suicide, post-traumatic stress disorder)<br>• History of neurological disorders (incl. multiple sclerosis, stroke, epilepsy, brain tumor, meningoencephalitis, severe concussion)<br>• History of malignant diseases<br>• Intake of one of the following medications:<br>• Any centrally active drugs (including Hypericum perforatum)<br>• Beta- and alpha-blocker<br>• Cortisol<br>• Any chemotherapeutic or psychopharmacological medication<br>• Positive drug anamnesis (extensive alcohol, MDMA, amphetamines, cocaine, opiates, benzodiazepine, cannabis)<br>• Body Mass Index < 18 or > 30<br>• Previous participation in any scientific study<br>• Past or present student of Psychology<br>• MRI exclusion criteria including:<br>• Any metallic implants, braces, non-removable piercings<br>• Tattoos<br>• Pregnancy<br>• Claustrophobia<br>• Tinnitus<br>• Surgical operation in the last 3 months |
| **Neural Consequences of Stress Study (sample 2) (Reinelt et al., in preparation)** | • Female sex<br>• Age < 20 or >35<br>• Smoking<br>• Past or present student of Psychology<br>• Excessive alcohol or drug consumption<br>• Regular medication intake<br>• History of cardiovascular or neurological diseases<br>• Body Mass Index > 27<br>• Positive drug anamnesis (extensive alcohol, MDMA, amphetamines, cocaine, opiates, benzodiazepine, cannabis)<br>• Positive diagnosis in psychiatric screening of axis I disorders<br>• Abnormalities in analysis of blood screening<br>• MRI exclusion criteria including:<br>• Any metallic implants, braces, non-removable piercings<br>• Tattoos<br>• Pregnancy<br>• Claustrophobia<br>• Tinnitus<br>• Surgical operation in the last 3 months |
| **Neuroanatomy and Connectivity Protocol<sup>12</sup> (sample 3)** | • Age < 20 or >75<br>• History of psychiatric diseases that required inpatient treatment for longer than 2 weeks within the last 10 years (e.g. psychosis, attempted suicide, post-traumatic stress disorder)<br>• History of neurological disorders (incl. multiple sclerosis, stroke, epilepsy, brain tumor, meningoencephalitis, severe concussion)<br>• History of malignant diseases<br>• Intake of one of the following medications:<br>• Any centrally active drugs (including Hypericum perforatum)<br>• Beta- and alpha-blocker<br>• Cortisol<br>• Any chemotherapeutic or psychopharmacological medication<br>• Positive drug anamnesis (extensive alcohol, MDMA, amphetamines, cocaine, opiates, benzodiazepine, cannabis)<br>• Body Mass Index <18 or >30<br>• Extensive testing experience at the Max-Planck-Institute or other academic institution<br>• Past or present student of Psychology<br>• MRI exclusion criteria:<br>• Any metallic implants, braces, non-removable piercings<br>• Tattoos<br>• Pregnancy<br>• Claustrophobia |
|  | • Tinnitus<br>• Surgical operation in the last 3 months |
| **Leipzig Research Centre for Civilization Diseases<sup>13</sup> (sample 4)** | • History of neurological disorders (incl. multiple sclerosis, stroke, epilepsy, parkinson's disease, brain tumor)<br>• History of malignant diseases<br>• History of depression<br>• History of cardiovascular disease (e.g. myoinfarction, coronary heart disease, heart surgery, bypass, catheter, stent)<br>• Hypertension<br>• Intake of anti-hypertensive drugs<br>• Intake of centrally-active drugs<br>• Diabetes (Type I or II)<br>• MRI exclusion criteria:<br>• Any metallic implants, braces, non-removable piercings<br>• Tattoos<br>• Pregnancy<br>• Claustrophobia<br>• Tinnitus<br>• Surgical operation in the last 3 months |

**Supplementary Table 2.**
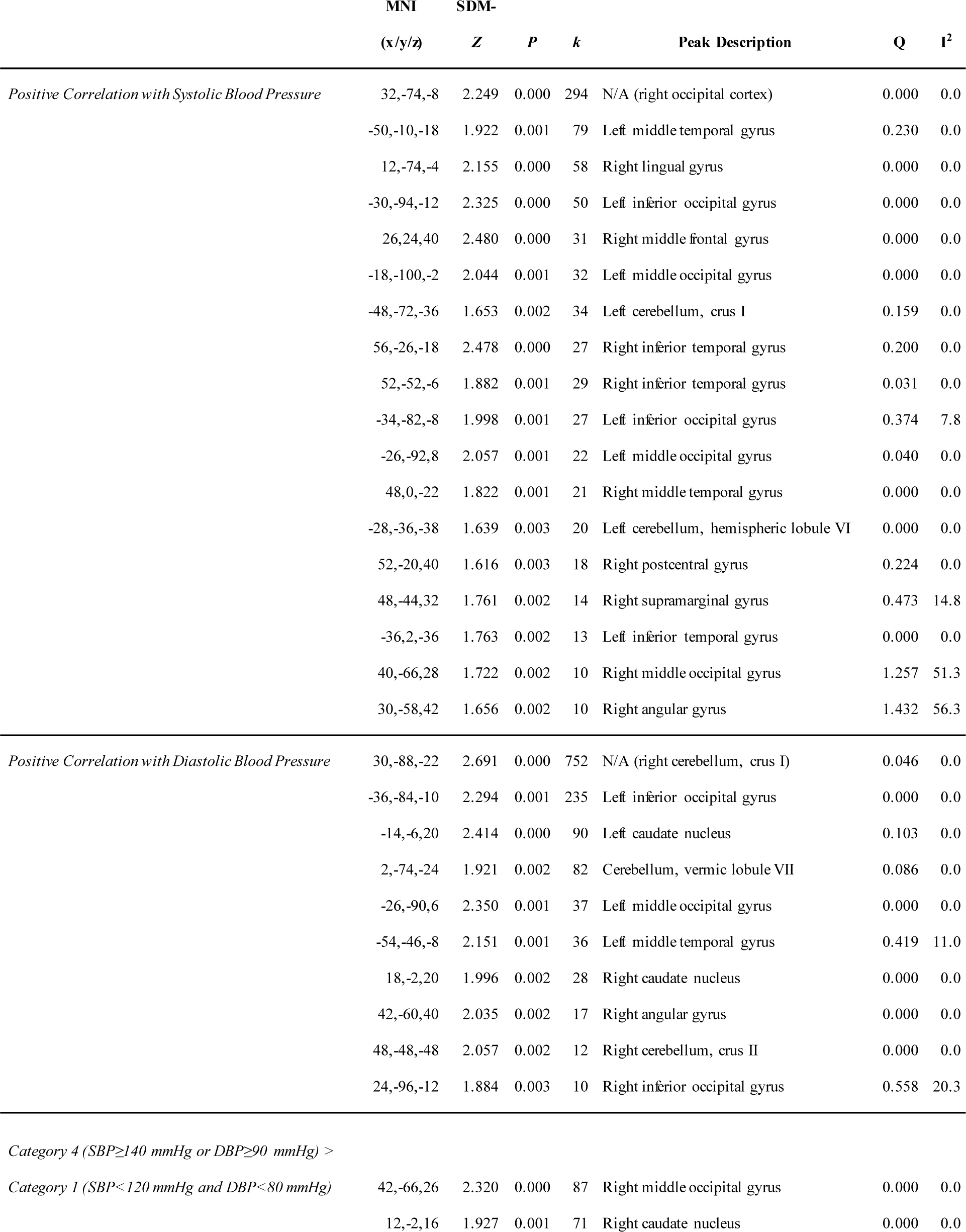
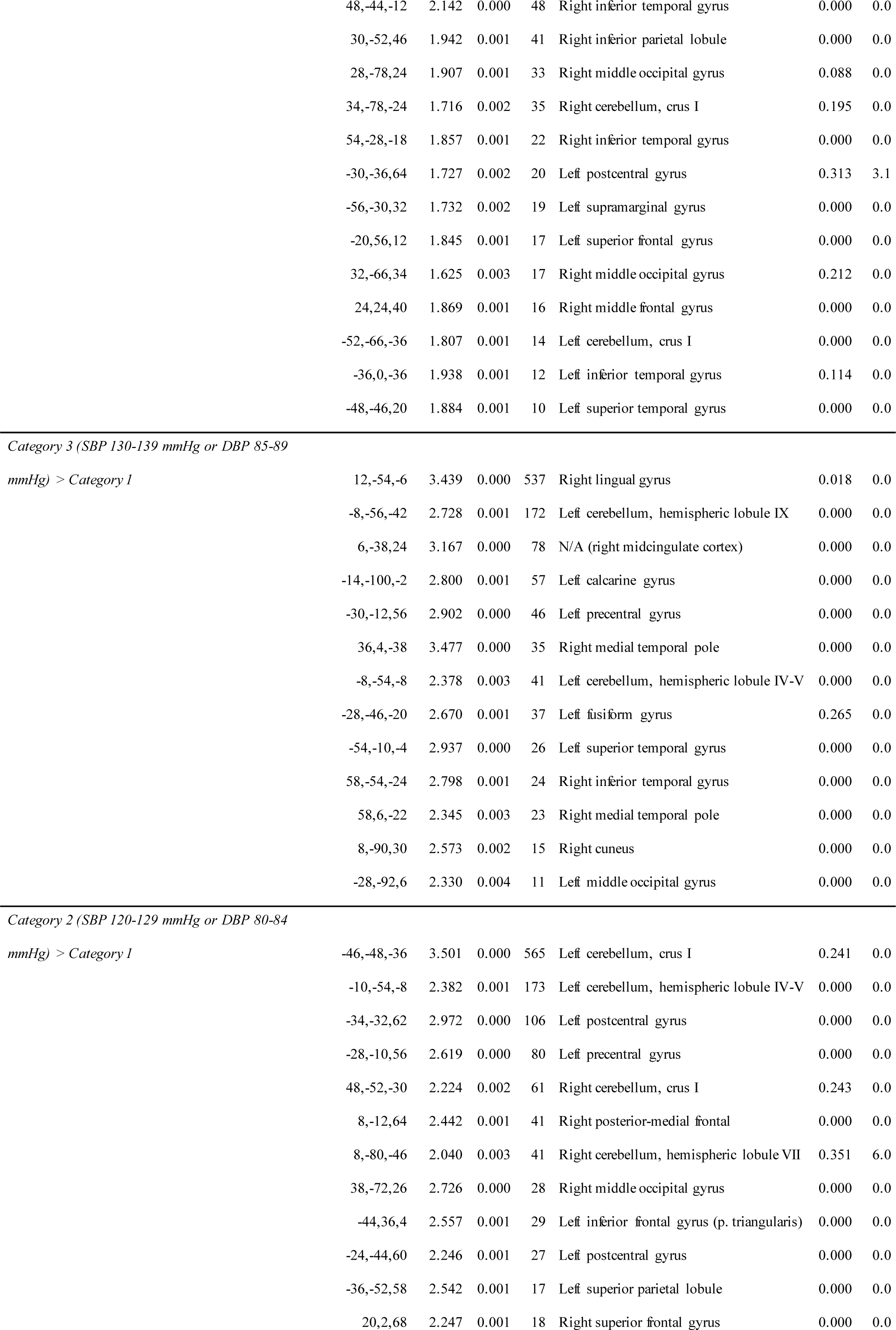
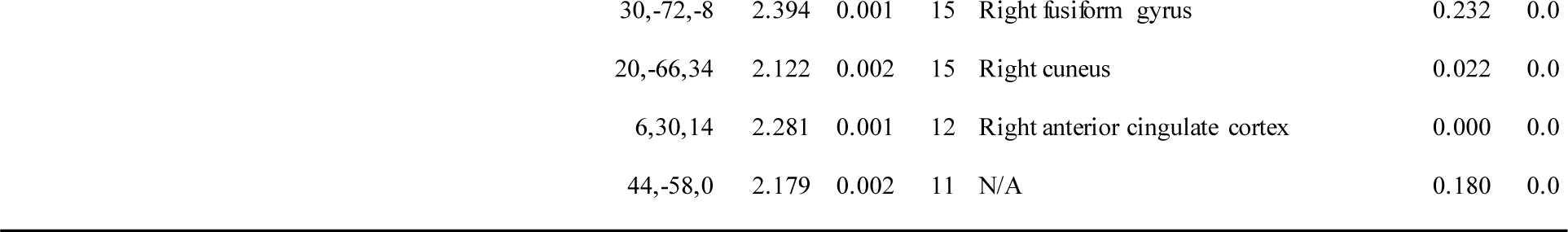
Positive image-based meta-analysis results of regional grey matter volume differences associated with blood pressure. Image-based meta-analysis results of significant clusters yielding higher grey matter volume for the respective contrast of interest. Columns indicate cluster-specific MNI coordinates of peak voxels, meta-analytic SDM-*Z*-value, meta-analytic *p*- value, number of voxels in cluster and anatomical label of the peak voxel. Anatomical labels were assigned using SPM’s Anatomy toolbox^8^. *Q* and *I*^*2*^ are measures of meta-analytic heterogeneity. Voxel threshold was set to *p*<0.005, peak height threshold was set to SDM -*Z*>1.0 and cluster extent threshold was set to *k*≥10 voxels as recommended by Radua et al. (2012)^7^. Final voxel size was 2 × 2 × 2 mm^3^. MNI: Montreal Neurological Institute. SDM: Seed-based *d* Mapping. SBP: Systolic blood pressure. DBP: Diastolic blood pressure.

**Supplementary Table 3.**
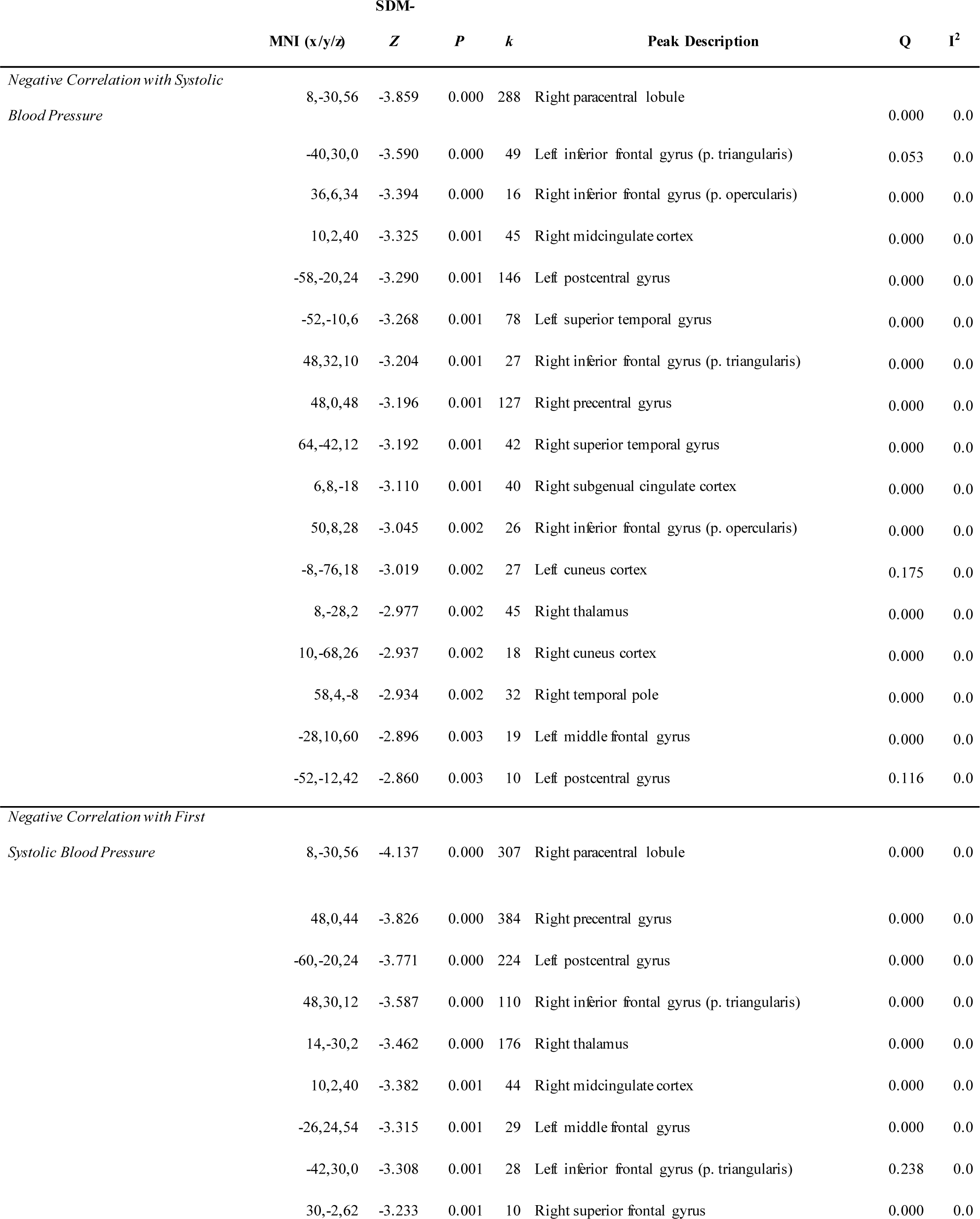
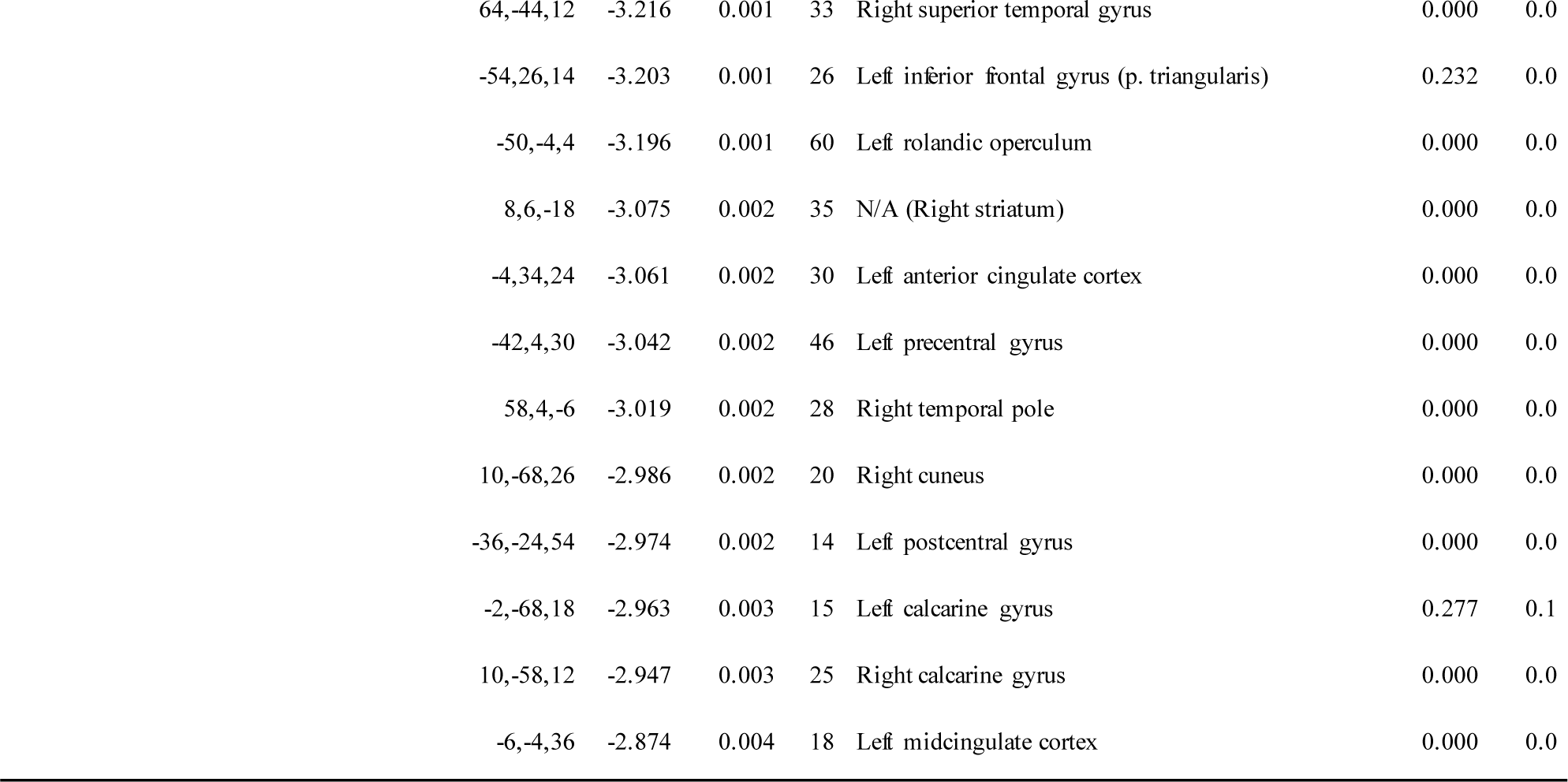
Image-based meta-analysis results of regional grey matter volume differences associated with first systolic blood pressure. Image-based meta-analysis results of significant clusters yielding lower grey matter volume for the respective contrast of interest (averaged SBP, first SBP). Columns indicate cluster-specific MNI coordinates of peak voxels, meta-analytic SDM-*Z*- value, meta-analytic *p*-value, number of voxels in cluster and anatomical label of the peak voxel. *Q*- and *I*^*2*^-statistics are estimates of meta-analytic heterogeneity of effects across studies. Anatomical labels were assigned using SPM’s Anatomy toolbox^8^. Voxel threshold was set to *p*<0.005, peak height threshold was set to SDM -*Z*>1.0 and cluster extent threshold was set to *k*≥10 voxels. Final voxel size was 2 × 2 × 2 mm^3^. MNI: Montreal Neurological Institute. SDM: Seed-based *d* Mapping.

| SDM- |  |  |  |  |  |  |  |
| --- | --- | --- | --- | --- | --- | --- | --- |
|  | MNI (x/y/z) | Z | P | k | Peak Description | Q | I<sup>2</sup> |
| Negative Correlation with Systolic | 8,-30,56 | -3.859 | 0.000 | 288 | Right paracentral lobule |  |  |
| Blood Pressure |  |  |  |  |  | 0.000 | 0.0 |
|  | -40,30,0 | -3.590 | 0.000 | 49 | Left inferior frontal gyrus (p. triangularis) | 0.053 | 0.0 |
|  | 36,6,34 | -3.394 | 0.000 | 16 | Right inferior frontal gyrus (p. opercularis) | 0.000 | 0.0 |
|  | 10,2,40 | -3.325 | 0.001 | 45 | Right midcingulate cortex | 0.000 | 0.0 |
|  | -58,-20,24 | -3.290 | 0.001 | 146 | Left postcentral gyrus | 0.000 | 0.0 |
|  | -52,-10,6 | -3.268 | 0.001 | 78 | Left superior temporal gyrus | 0.000 | 0.0 |
|  | 48,32,10 | -3.204 | 0.001 | 27 | Right inferior frontal gyrus (p. triangularis) | 0.000 | 0.0 |
|  | 48,0,48 | -3.196 | 0.001 | 127 | Right precentral gyrus | 0.000 | 0.0 |
|  | 64,-42,12 | -3.192 | 0.001 | 42 | Right superior temporal gyrus | 0.000 | 0.0 |
|  | 6,8,-18 | -3.110 | 0.001 | 40 | Right subgenual cingulate cortex | 0.000 | 0.0 |
|  | 50,8,28 | -3.045 | 0.002 | 26 | Right inferior frontal gyrus (p. opercularis) | 0.000 | 0.0 |
|  | -8,-76,18 | -3.019 | 0.002 | 27 | Left cuneus cortex | 0.175 | 0.0 |
|  | 8,-28,2 | -2.977 | 0.002 | 45 | Right thalamus | 0.000 | 0.0 |
|  | 10,-68,26 | -2.937 | 0.002 | 18 | Right cuneus cortex | 0.000 | 0.0 |
|  | 58,4,-8 | -2.934 | 0.002 | 32 | Right temporal pole | 0.000 | 0.0 |
|  | -28,10,60 | -2.896 | 0.003 | 19 | Left middle frontal gyrus | 0.000 | 0.0 |
|  | -52,-12,42 | -2.860 | 0.003 | 10 | Left postcentral gyrus | 0.116 | 0.0 |
| Negative Correlation with First |  |  |  |  |  |  |  |
| Systolic Blood Pressure | 8,-30,56 | -4.137 | 0.000 | 307 | Right paracentral lobule | 0.000 | 0.0 |
|  | 48,0,44 | -3.826 | 0.000 | 384 | Right precentral gyrus | 0.000 | 0.0 |
|  | -60,-20,24 | -3.771 | 0.000 | 224 | Left postcentral gyrus | 0.000 | 0.0 |
|  | 48,30,12 | -3.587 | 0.000 | 110 | Right inferior frontal gyrus (p. triangularis) | 0.000 | 0.0 |
|  | 14,-30,2 | -3.462 | 0.000 | 176 | Right thalamus | 0.000 | 0.0 |
|  | 10,2,40 | -3.382 | 0.001 | 44 | Right midcingulate cortex | 0.000 | 0.0 |
|  | -26,24,54 | -3.315 | 0.001 | 29 | Left middle frontal gyrus | 0.000 | 0.0 |
|  | -42,30,0 | -3.308 | 0.001 | 28 | Left inferior frontal gyrus (p. triangularis) | 0.238 | 0.0 |
|  | 30,-2,62 | -3.233 | 0.001 | 10 | Right superior frontal gyrus | 0.000 | 0.0 |

|  |  |  |  |  |  |  |
| --- | --- | --- | --- | --- | --- | --- |
| 64,-44,12 | -3.216 | 0.001 | 33 | Right superior temporal gyrus | 0.000 | 0.0 |
| -54,26,14 | -3.203 | 0.001 | 26 | Left inferior frontal gyrus (p. triangularis) | 0.232 | 0.0 |
| -50,-4,4 | -3.196 | 0.001 | 60 | Left rolandic operculum | 0.000 | 0.0 |
| 8,6,-18 | -3.075 | 0.002 | 35 | N/A (Right striatum) | 0.000 | 0.0 |
| -4,34,24 | -3.061 | 0.002 | 30 | Left anterior cingulate cortex | 0.000 | 0.0 |
| -42,4,30 | -3.042 | 0.002 | 46 | Left precentral gyrus | 0.000 | 0.0 |
| 58,4,-6 | -3.019 | 0.002 | 28 | Right temporal pole | 0.000 | 0.0 |
| 10,-68,26 | -2.986 | 0.002 | 20 | Right cuneus | 0.000 | 0.0 |
| -36,-24,54 | -2.974 | 0.002 | 14 | Left postcentral gyrus | 0.000 | 0.0 |
| -2,-68,18 | -2.963 | 0.003 | 15 | Left calcarine gyrus | 0.277 | 0.1 |
| 10,-58,12 | -2.947 | 0.003 | 25 | Right calcarine gyrus | 0.000 | 0.0 |
| -6,-4,36 | -2.874 | 0.004 | 18 | Left midcingulate cortex | 0.000 | 0.0 |

## Supplementary Figures

**Supplementary Figure 1.**
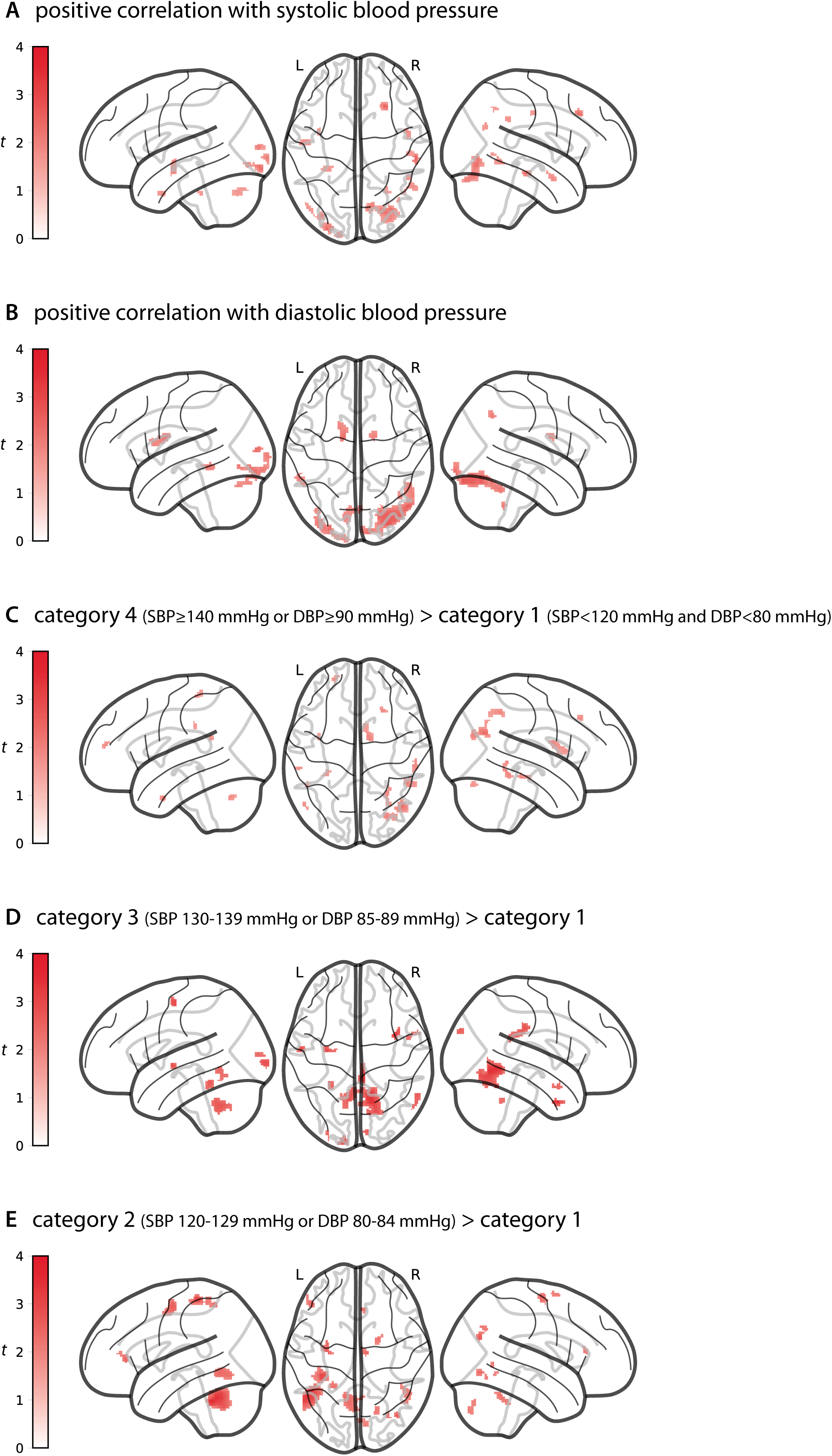
Meta-analytic positive differences in grey matter volume between blood pressure categories. Glass brain views of image-based meta-analysis results for the blood pressure category contrasts of interest with relevant slice views below (**A-E**). **A** and **B** depict positive correlations between SBP/DBP and gray matter volume, respectively. Red clusters indicate meta-analytic grey matter volume differences for the given contrast at a voxel threshold of *p*<0.005 with peak height threshold of SDM -*Z*>1.0 and cluster extent threshold of *k*≥10 (validated for high meta-analytic sensitivity and specificity (Radua et al., 2012)^7^). Color bars represent SDM -*Z*-values. 3D-volumetric results of these analyses can be inspected in detail on http://neurovault.org/collections/FDWHFSYZ/. SDM: Seed-based *d* Mapping. SBP: Systolic blood pressure. DBP: Diastolic blood pressure. L: Left hemisphere. R: Right hemisphere.

**Supplementary Figure 2.**
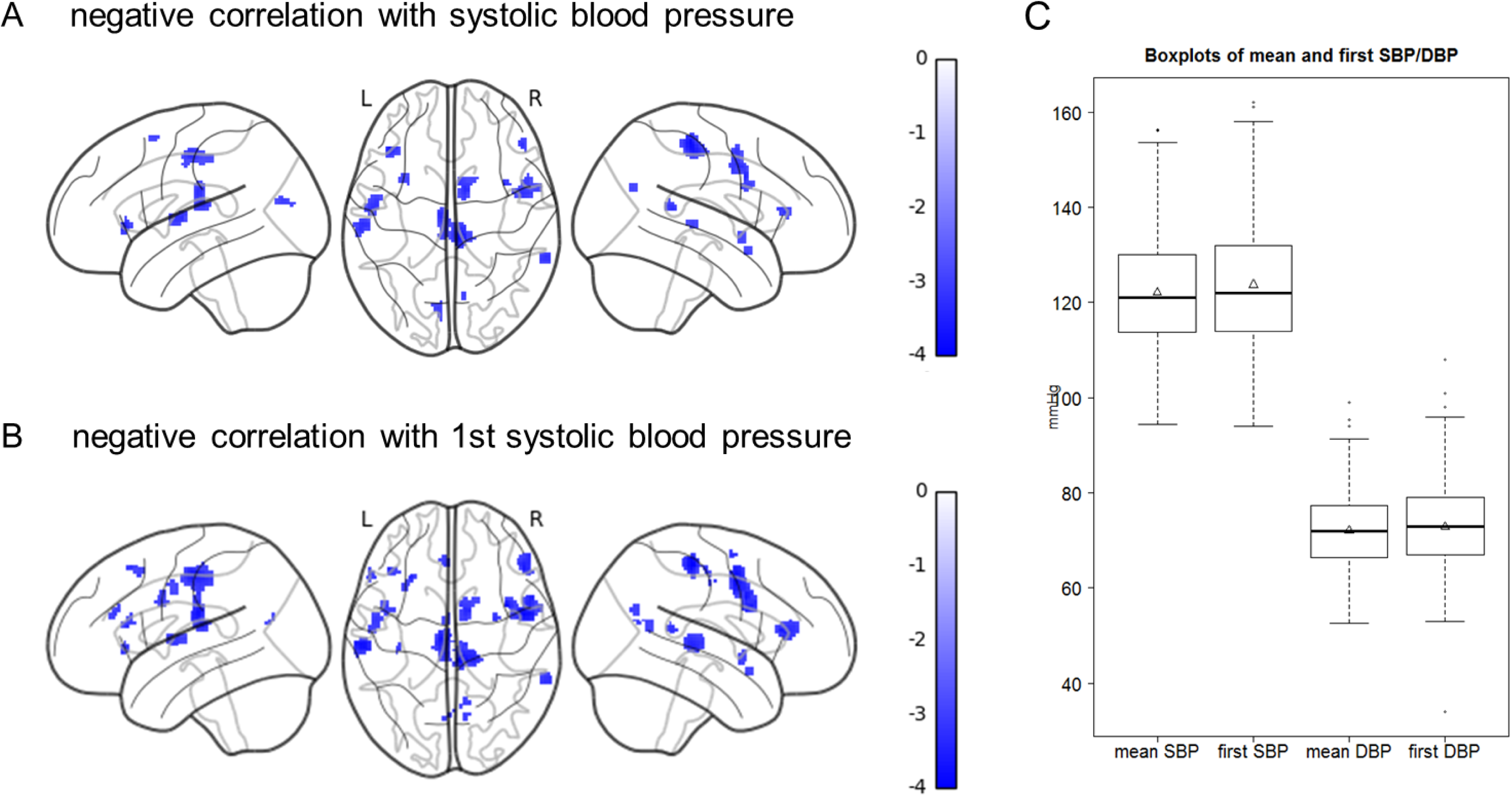
Associations between grey matter volume and first systolic BP reading A: Glass brain views of image-based meta-analysis results for the negative correlation between systolic blood pressure and gray matter volume in all four samples (n=423). **B**: Glass brain views of image-based meta-analysis results for the negative correlation between first measured systolic blood pressure and gray matter volume in all four samples. Blue clusters indicate meta-analytic gray matter volume differences at a voxel threshold of *p*<0.005 with peak height threshold of SDM -*Z*<-1.0 and cluster extent threshold of *k*≥10. Color bars represent SDM -*Z*-values. C: Boxplots of mean and first SBP or DBP, respectively, in study samples with ≥2 BP readings (samples 1, 2 and 4, n=353). Triangles represent the mean. SDM: Seed-based *d* Mapping. SBP: Systolic blood pressure. DBP: Diastolic blood pressure. L: Left hemisphere. R: Right hemisphere.

Author contributions
*Study concept and design:* Schaare, Villringer.
*Statistical analysis:* Schaare.
*Acquisition or interpretation of data:* All authors.
*Drafting of the manuscript:* Schaare, Villringer.
*Critical revision of the manuscript:* All authors.

## Acknowledgements

We thank all volunteers for their participation in any of the studies. Furthermore, we thank all researchers, technicians and students who planned, collected, entered and curated data used in this manuscript.

## References

1. Forouzanfar MH, Afshin A, Alexander LT, et al. Global, regional, and national comparative risk assessment of 79 behavioural, environmental and occupational, and metabolic risks or clusters of risks, 1990–2015: a systematic analysis for the Global Burden of Disease Study 2015. Lancet. 2016;388(10053):1659–1724. doi:10.1016/S0140-6736(16)31679-8

2. NCD Risk Factor Collaboration (NCD-RisC). Worldwide trends in blood pressure from 1975 to 2015: a pooled analysis of 1479 population-based measurement studies with 19·1 million participants. Lancet. 2016;389(10064):634–647. doi:10.1016/S0140-6736(16)31919-5

3. Iadecola C, Yaffe K, Biller J, et al. Impact of hypertension on cognitive function?: a scientific statement from the American Heart Association. Hypertension. 2016;68:e67– e94. doi:10.1161/HYP.0000000000000053

4. Norton S, Matthews FE, Barnes DE, Yaffe K, Brayne C. Potential for primary prevention of Alzheimer’s disease: An analysis of population-based data. Lancet Neurol. 2014;13(8):788–794. doi:10.1016/S1474-4422(14)70136-X

5. Power MC, Schneider ALC, Wruck L, et al. Life-course blood pressure in relation to brain volumes. Alzheimer’s Dement. 2016;12(8):890–899. doi:10.1016/j.jalz.2016.03.012

6. Hajjar I, Zhao P, Alsop D, et al. Association of blood pressure elevation and nocturnal dipping with brain atrophy, perfusion and functional measures in stroke and nonstroke individuals. Am J Hypertens. 2010;23(1):17–23. doi:10.1038/ajh.2009.187

7. Launer LJ, Lewis CE, Schreiner PJ, et al. Vascular factors and multiple measures of early brain health: CARDIA brain MRI study. Ikram MA, ed. PLoS One. 2015;10(3):e0122138. doi:10.1371/journal.pone.0122138

8. Petrovitch H, White LR, Izmirilian G, et al. Midlife blood pressure and neuritic plaques, neurofibrillary tangles, and brain weight at death: the HAAS. Neurobiol Aging. 2000;21(1):57–62. doi:10.1016/S0197-4580(00)00106-8

9. den Heijer T, Launer LJ, Prins ND, et al. Association between blood pressure, white matter lesions, and atrophy of the medial temporal lobe. Neurology. 2005;64(2):263–267. doi:10.1212/01.WNL.0000149641.55751.2E

10. Raz N, Lindenberger U, Rodrigue KM, et al. Regional brain changes in aging healthy adults: general trends, individual differences and modifiers. Cereb Cortex. 2005;15(11):1676–1689. doi:10.1093/cercor/bhi044

11. Leritz EC, Salat DH, Williams VJ, et al. Thickness of the human cerebral cortex is associated with metrics of cerebrovascular health in a normative sample of community dwelling older adults. Neuroimage. 2011;54(4):2659–2671. doi:10.1016/j.neuroimage.2010.10.050

12. Debette S, Seshadri S, Beiser A, et al. Midlife vascular risk factor exposure accelerates structural brain aging and cognitive decline. Neurology. 2011;77(5):461–468. doi:10.1212/WNL.0b013e318227b227

13. Maillard P, Seshadri S, Beiser A, et al. Effects of systolic blood pressure on white-matter integrity in young adults in the Framingham Heart Study: a cross-sectional study. Lancet Neurol. 2012;11(12):1039–1047. doi:10.1016/S1474-4422(12)70241-7

14. Muller M, Sigurdsson S, Kjartansson O, et al. Joint effect of mid-and late-life blood pressure on the brain: the AGES-Reykjavik study. Neurology. 2014;82(24):2187–2195. doi:10.1212/WNL.0000000000000517

15. Beauchet O, Celle S, Roche F, et al. Blood pressure levels and brain volume reduction: a systematic review and meta-analysis. J Hypertens. 2013;31(8):1502–1516. doi:10.1097/HJH.0b013e32836184b5

16. Olsen MH, Angell SY, Asma S, et al. A call to action and a lifecourse strategy to address the global burden of raised blood pressure on current and future generations: the Lancet Commission on hypertension. Lancet. 2016;388(10060):2665–2712. doi:10.1016/S0140-6736(16)31134-5

17. Grossman DC, Bibbins-Domingo K, Curry SJ, et al. Behavioral Counseling to Promote a Healthful Diet and Physical Activity for Cardiovascular Disease Prevention in Adults Without Cardiovascular Risk Factors. JAMA. 2017;318(2):167. doi:10.1001/jama.2017.7171

18. Gianaros PJ, Sheu LK, Matthews K a, Jennings JR, Manuck SB, Hariri AR. Individual differences in stressor-evoked blood pressure reactivity vary with activation, volume, and functional connectivity of the amygdala. J Neurosci. 2008;28(4):990–999. doi:10.1523/JNEUROSCI.3606-07.2008

19. Ashburner J, Friston KJ. Voxel-based morphometry--the methods. Neuroimage. 2000;11(6 Pt 1):805–821. doi:10.1006/nimg.2000.0582

20. Ashburner J. A fast diffeomorphic image registration algorithm. Neuroimage. 2007;38(1):95–113. doi:10.1016/j.neuroimage.2007.07.007

21. Mendes N, Oligschlaeger S, Lauckner ME, et al. A functional connectome phenotyping dataset including cognitive state and personality measures. bioRxiv. 2017. http://www.biorxiv.org/content/early/2017/07/18/164764. Accessed July 19, 2017.

22. Loeffler M, Engel C, Ahnert P, et al. The LIFE-Adult-Study: objectives and design of a population-based cohort study with 10,000 deeply phenotyped adults in Germany. BMC Public Health. 2015;15:691. doi:10.1186/s12889-015-1983-z

23. Mancia G, Fagard R, Narkiewicz K, et al. 2013 ESH/ESC Guidelines for the management of arterial hypertension. Eur Heart J. 2013;34(28):2159–2219. doi:10.1093/eurheartj/eht151

24. Radua J, Mataix-Cols D, Phillips ML, et al. A new meta-analytic method for neuroimaging studies that combines reported peak coordinates and statistical parametric maps. Eur Psychiatry. 2012;27(8):605–611. doi:10.1016/j.eurpsy.2011.04.001

25. Fazekas F, Chawluk J, Alavi A, Hurtig H, Zimmerman R. MR signal abnormalities at 1.5 T in Alzheimer’s dementia and normal aging. Am J Roentgenol. 1987;149(2):351–356. doi:10.2214/ajr.149.2.351

26. Diedrichsen J. A spatially unbiased atlas template of the human cerebellum. Neuroimage. 2006;33(1):127–138. doi:10.1016/j.neuroimage.2006.05.056

27. Critchley HD, Harrison N a. Visceral influences on brain and behavior. Neuron. 2013;77(4):624–638. doi:10.1016/j.neuron.2013.02.008

28. Woo MA, Macey PM, Fonarow GC, Hamilton MA, Harper RM. Regional brain gray matter loss in heart failure. J Appl Physiol. 2003;95(2):677–684. doi:10.1152/japplphysiol.00101.2003

29. Avelar WM, D’Abreu A, Coan AC, et al. Asymptomatic carotid stenosis is associated with gray and white matter damage. Int J Stroke. 2015;10(8):1197–1203. doi:10.1111/ijs.12574

30. Lorio S, Lutti A, Kherif F, et al. Disentangling in vivo the effects of iron content and atrophy on the ageing human brain. Neuroimage. 2014;103:280–289. doi:10.1016/j.neuroimage.2014.09.044

31. Critchley HD, Mathias CJ, Josephs O, et al. Human cingulate cortex and autonomic control: Converging neuroimaging and clinical evidence. Brain. 2003;126(10):2139–2152. doi:10.1093/brain/awg216

32. Oppenheimer SM, Kedem G, Martin WM. Left-insular cortex lesions perturb cardiac autonomic tone in humans. Clin Auton Res. 1996;6(3):131–140. doi:10.1007/BF02281899

33. Krause T, Werner K, Fiebach JB, et al. Stroke in right dorsal anterior insular cortex Is related to myocardial injury. Ann Neurol. 2017;81(4):502–511. doi:10.1002/ana.24906

34. Braak H, Braak E. Neuropathological stageing of Alzheimer-related changes. Acta Neuropathol. 1991;82:239–259. doi:10.1007/BF00308809

35. Dickerson BC, Stoub TR, Shah RC, et al. Alzheimer-signature MRI biomarker predicts AD dementia in cognitively normal adults. Neurology. 2011;76(16):1395–1402. doi:10.1212/WNL.0b013e3182166e96

36. Allan C, Zsoldos E, Filippini N, et al. Life-time hypertension as a predictor of brain structure in older adults: a prospective cohort study. Br J Psychiatry. 2015;206(4):308–315. doi:10.1192/bjp.bp.114.153536

37. Muller M, van der Graaf Y, Visseren FL, Vlek AL, Mali WPt, Geerlings MI. Blood pressure, cerebral blood flow, and brain volumes. The SMART-MR study. J Hypertens. 2010;28(7):1498–1505. doi:10.1097/HJH.0b013e32833951ef

38. Kennedy KM, Erickson KI, Rodrigue KM, et al. Age-related differences in regional brain volumes: A comparison of optimized voxel-based morphometry to manual volumetry. Neurobiol Aging. 2009;30(10):1657–1676. doi:10.1016/j.neurobiolaging.2007.12.020

39. Tardif CL, Steele CJ, Lampe L, et al. Investigation of the confounding effects of vasculature and metabolism on computational anatomy studies. Neuroimage. 2017;149:233–243. doi:10.1016/j.neuroimage.2017.01.025

40. Strong JP, Malcom GT, Mcmahan CA, et al. Prevalence and Extent of Atherosclerosis in Adolescents and Young Adults. JAMA. 1999;281(8):727–735. doi:10.1001/jama.281.8.727

41. Maillard P, Mitchell GF, Himali JJ, et al. Effects of arterial stiffness on brain integrity in young adults from the framingham heart study. Stroke. 2016;47(4):1030–1036. doi:10.1161/STROKEAHA.116.012949

42. Neuhauser HK, Adler C, Rosario AS, Diederichs C, Ellert U. Hypertension prevalence, awareness, treatment and control in Germany 1998 and 2008-11. J Hum Hypertens. 2015;29(August):1–7. doi:10.1038/jhh.2014.82

43. Chobanian A V, Bakris GL, Black HR, et al. Seventh report of the Joint National Committee on Prevention, Detection, Evaluation, and Treatment of High Blood Pressure. Hypertension. 2003;42(6):1206–1252. doi:10.1161/01.HYP.0000107251.49515.c2

44. Whelton PK, Carey RM, Aronow WS, et al. 2017 ACC/AHA/AAPA/ABC/ACPM/AGS/APhA/ASH/ASPC/NMA/PCNA Guideline for the Prevention, Detection, Evaluation, and Management of High Blood Pressure in Adults. Hypertension. November 2017:HYP.0000000000000065. doi:10.1161/HYP.0000000000000065

## References

1. Marques JP, Kober T, Krueger G, van der Zwaag W, Van de Moortele P-F, Gruetter R. MP2RAGE, a self bias-field corrected sequence for improved segmentation and T1-mapping at high field. Neuroimage. 2010;49(2):1271–1281. doi:10.1016/j.neuroimage.2009.10.002

2. Streitbürger D-P, Pampel A, Krueger G, et al. Impact of image acquisition on voxel-based-morphometry investigations of age-related structural brain changes. Neuroimage. 2014;87:170–182. doi:10.1016/ j.neuroimage.2013.10.051

3. Jenkinson M, Beckmann CF, Behrens TEJ, Woolrich MW, Smith SM. FSL. Neuroimage. 2012;62(2):782–790. doi:10.1016/ j.neuroimage.2011.09.015

4. Ashburner J, Friston KJ. Voxel-based morphometry--the methods. Neuroimage. 2000;11(6 Pt 1):805–821. doi:10.1006/nimg.2000.0582

5. Ashburner J. A fast diffeomorphic image registration algorithm. Neuroimage. 2007;38(1):95–113. doi:10.1016/ j.neuroimage.2007.07.007

6. Gorgolewski KJ, Varoquaux G, Rivera G, et al. NeuroVault.org: a web-based repository for collecting and sharing unthresholded statisticalmaps of the human brain. Front Neuroinform. 2015;9:8. doi:10.3389/fninf.2015.00008

7. Radua J, Mataix-Cols D, Phillips ML, et al. A new meta-analytic method for neuroimaging studies that combines reported peak coordinates and statistical parametric maps. Eur Psychiatry. 2012;27(8):605–611. doi:10.1016/j.eurpsy.2011.04.001

8. Eickhoff SB, Paus T, Caspers S, et al. Assignment of functional activations to probabilistic cytoarchitectonic areas revisited. Neuroimage. 2007;36(3):511–521. doi:10.1016/ j.neuroimage.2007.03.060

9. Abraham A, Pedregosa F, Eickenberg M, et al. Machine learning for neuroimaging with scikit-learn. Front Neuroinform. 2014;8(February):14. doi:10.3389/fninf.2014.00014

10. Pickering TG, Hall JE, Appel LJ, et al. Recommendations for blood pressure measurement in humans and experimental animals: Part 1: blood pressure measurement in humans: a statement for professionals from the Subcommittee of Professional and Public Education of the American Heart Association Cou. Hypertens. 2005;45(1):142–161. doi:10.1161/01.HYP.0000150859.47929.8e

11. Higgins JPT, Green S (eds.). Cochrane Handbook for Systematic Reviews of Interventions Version 5.1.0 [updated March 2011]. The Cochrane Collaboration, 2011. Available from http://handbook.cochrane.org.

12. Mendes N, Oligschlaeger S, Lauckner ME, et al. A functional connectome phenotyping dataset including cognitive state and personality measures. bioRxiv. 2017. http://www.biorxiv.org/content/early/2017/07/18/164764. Accessed July 19, 2017.

13. Loeffler M, Engel C, Ahnert P, et al. The LIFE-Adult-Study: objectives and design of a population-based cohort study with 10,000 deeply phenotyped adults in Germany. BMC Public Health. 2015;15:691. doi:10.1186/s12889-015-1983-z

